# RNAlight: a machine learning model to identify nucleotide features determining RNA subcellular localization

**DOI:** 10.1101/2022.09.16.508211

**Authors:** Guo-Hua Yuan, Ying Wang, Guang-Zhong Wang, Li Yang

## Abstract

Different RNAs have distinct subcellular localizations. However, nucleotide features that determine these distinct distributions of lncRNAs and mRNAs have yet to be fully addressed. Here, we develop RNAlight, a machine learning model based on LightGBM, to identify nucleotide *k*-mers contributing to the subcellular localizations of mRNAs and lncRNAs. With the Tree SHAP algorithm, RNAlight extracts nucleotide features for cytoplasmic or nuclear localization of RNAs, indicating the sequence basis for distinct RNA subcellular localizations. By assembling *k*-mers to sequence features and subsequently mapping to known RBP-associated motifs, different types of sequence features and their associated RBPs were additionally uncovered for lncRNAs and mRNAs with distinct subcellular localizations. Finally, we extended RNAlight to precisely predict the subcellular localizations of other types of RNAs, including snRNAs, snoRNAs and different circular RNA transcripts, suggesting the generality of using RNAlight for RNA subcellular localization prediction.

**Key points:**

- A machine learning model, RNAlight, is developed to efficiently and sensitively predict subcellular localizations of mRNAs and lncRNAs.
- With embedded Tree SHAP algorithm, RNAlight further reveals distinct key sequence features and their associated RBPs for subcellular localizations of mRNAs or lncRNAs.
- RNAlight is successfully extended for the subcellular localization prediction of additional types of noncoding RNAs that were not used for model development, such as circular RNAs, suggesting its generality in RNA subcellular localization prediction.
- RNAlight is freely available at https://github.com/YangLab/RNAlight.

## Introduction

RNA localization is closely related to its biogenesis, processing and function, which also determines cell fate and polarity [1, 2]. In general, most messenger RNAs (mRNAs) transcribed from protein-coding gene loci are usually processed with a series of co- and/or post-transcriptional regulation, including but not limited to 5’-cap, splicing, editing/modification and 3’-adenylation, and transported from nucleus to cytoplasm for protein translation [3, 4]. Instead, many well-studied long non-coding RNAs (lncRNAs) with the length of more than 200 nucleotides tend to be located in nucleus to regulate gene expression by associating with chromatin [5]. Nevertheless, a set of mRNA transcripts can be temporarily retained in nucleus, possibly due to the existence of inverted repeated elements in their 3’ untranslated regions (3’ UTR), by which the translation of specific proteins is retarded [6–9]. Interestingly, some lncRNAs can be exported to the cytoplasm to regulate protein translation by associating with miRNA or ribosome [10, 11]. Thus, understanding RNA molecules’ subcellular localizations is important to their functional study.

A variety of approaches have been applied to study RNA subcellular localization. RNA fluorescence in situ hybridization (FISH) can accurately examine RNA subcellular localization in a single-RNA resolution and in living cells [12–14]. In addition, cytoplasmic and nuclear RNAs can be biochemically separated into different proportions and further examined by RT-PCR for individual RNAs or by high-throughput methods for various RNA species on a genomewide scale. For example, CeFra-seq [15] extracted cell fractions of cytosol, insolubles, membrane and nucleus for high-throughput sequencing to identify RNA localization in these diverse cell fractions. Moreover, APEX-Seq [16], which is also an RNA-sequencing based method to examine direct proximity labeling of RNA using the peroxidase enzyme APEX2, revealed extensive patterns of localization for diverse RNA classes in distinct subcellular locales. Furthermore, by collecting the subcellular localization information of thousands of RNAs across different cell lines and species, multiple databases, such as LncATLAS [17] and RNALocate [18], have been constructed to summarize RNA subcellular localization on a genome-wide scale. These datasets not only provide information of individual RNA subcellular localization, but also render a foundation for the prediction of RNA subcellular location *in silico.* In this case, several machine learning and deep learning methods, including mRNALoc [19] and DeepLncRNA [20], have been established to predict RNA subcellular localization. Specifically, the mRNALoc model examined cytoplasmic and nuclear localizations of mRNAs by support vector machine (SVM) based on sequence information [19], while the DeepLncRNA pipeline [20] trained a deep learning framework to predict nucleus to cytoplasm ratios of lncRNAs. Although these models have been successfully used to predict the localization of RNA based on sequence information, it remained unclear what kinds of key sequence features may contribute to the distinct localization of different types of RNAs. Meanwhile, since these reported methods were applicable to the prediction for just one specific type of RNA, mRNA or lncRNA [19–23], a universal model to perform subcellular localization prediction for different types of RNAs has been lacking.

In this study, we developed a machine learning model, RNAlight, which is based on Light Gradient Boosting Machine (LightGBM) [24], to simultaneously predict the subcellular localizations of both mRNA and lncRNA by using *k*-mer frequencies as input. With the integration of Shapley Additive exPlanations (Tree SHAP) [25] and *k*-mer assembly in RNAlight, different sequence features and their associated RNA binding proteins (RBPs) that contribute to distinct subcellular localizations of mRNA or lncRNA were further revealed. With RNAlight, subcellular localizations of various types of noncoding RNAs, including snRNAs, snoRNAs and circular RNAs, were also predicted in line with their reported functions, suggesting its general application in the study of RNA localization and function.

## Results

### Training RNAlight model to predict RNA subcellular localization

To train models for the prediction of RNA subcellular localizations, three published datasets, including CeFra-seq [15], APEX-Seq [16] and LncATLAS [17] (Figure 1A; Supplementary Figure S1A-B), were collected to construct a combined RNA subcellular localization library for both lncRNAs (n = 4,623) and mRNAs (n = 6,245) with GENCODE v30 annotation file (http://ftp.ebi.ac.uk/pub/databases/gencode/Gencode_human/release_30/gencode.v30.annotation.gtf.gz). Of note, only two major subcellular localizations, nucleus and cytoplasm, were used for model training and prediction in this study. Among four fractions in the CeFra-seq dataset, RNAs in the cytosol, insoluble and membrane fractions from the general cytoplasmic extract were all considered as cytoplasmic localization, and ones in the nuclear fraction were considered as nuclear localization. Among eight classifications in the APEX-Seq dataset, cytoplasm, ER membrane, ER lumen and outer mitochondrial membrane classifications were considered as the cytoplasm classification, while nucleus, nucleolus, lamina and nuclear pore classifications were considered as the nucleus classification. LncATLAS dataset classified lncRNAs with nuclear or cytoplasmic subcellular localization. Since experimentally examined in different conditions, such as various cell lines/methods, some RNAs were shown inconsistent with multiple localizations across these publicly-available datasets, and were removed from further processing. After this filter step, about 3,792 lncRNAs (1,986 nuclear and 1,806 cytoplasmic lncRNAs, Supplementary Table S1) and 5,180 mRNAs (2,256 nuclear and cytoplasmic 2,924 mRNAs, Supplementary Table S2) with consistent and single localization across different datasets were combined, and randomly split into training sets (Training-lncRNA, n = 3,412; Training-mRNA, n = 4,662) and test sets (Test-lncRNA, n = 380; Test-mRNA, n = 518) with a 9:1 ratio for model construction and evaluation, respectively (Supplementary Figure S1A-B) (“Methods” section).

**Figure 1.**
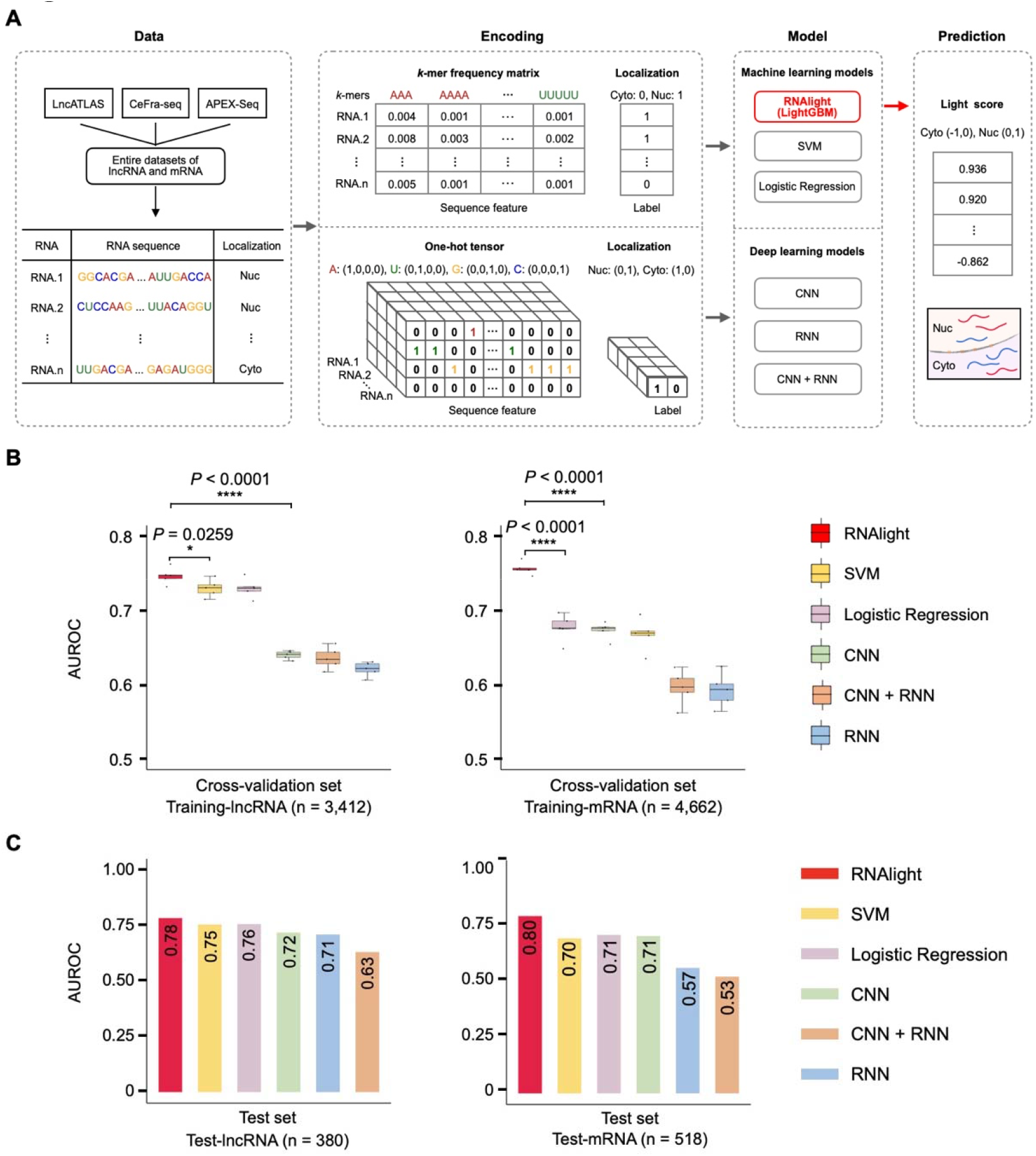
Computational models to predict RNA subcellular localization. (**A**) Schematic diagram of computational models for RNA subcellular localization prediction. (**B**) Crossvalidation of different models for lncRNA (left) and mRNA (right) subcellular localization prediction. Each dot represents the area under the receiver operating characteristic curve (AUROC) from five-fold cross-validation (total AUROC, n = 5). Statistical testing was performed with one-sided Welch’s *t*-test. In the box plots, the 25th, 50th and 75th percentiles are indicated as the top, middle, and bottom lines, respectively; whiskers represent the 10th and 90th percentiles, respectively. (**C**) Evaluation of prediction models for lncRNA (left) and mRNA (right) subcellular localization by using the test sets.

Next, we constructed a series of machine learning and deep learning models for the prediction of RNA subcellular localization (Figure 1A). These include three machine learning models, such as canonical support vector machine (SVM), logistic regression and an RNAlight model based on the LightGBM framework that uses tree-based learning algorithms by Microsoft [24], and three deep learning models, such as convolutional neural network (CNN), recurrent neural network (RNN) and a hybrid of CNN and RNN (CNN+RNN) (Figure 1A, Supplementary Figure S2A).

Specifically, we adopted corresponding featurization methods for machine learning [26] and deep learning[27] models, respectively. For machine learning models, RNA sequences were converted to the *k*-mer (*k* equals to 3, 4, or 5) frequency matrix as input features (Figure 1A, “Methods” section). For deep learning models, given that the classic CNN model only accepts the fixed-length input that is connected to the fully connected layer for classification or regression tasks [28–30], each RNA sequence was processed to a fixed length (lncRNA, 4,000nt; mRNA, 9,000nt) by padding or truncating, and further converted to the tensor by one-hot encoding as input (“Methods” section).

After training with the same sets (“Methods” section), we compared their performances and found that RNAlight showed the best performance in the prediction of RNA (both lncRNA and mRNA) localization with cross-validation, indicated by area under the receiver operating characteristic curve (AUROC) (Figure 1B). Consistently, when evaluated with test sets, RNAlight also outperformed other models with the AUROC values as 0.78 and 0.80 for predicting lncRNA or mRNA subcellular localization, respectively (Figure 1C; Table 1–2). Of note, all the deep learning models in our study have relatively poor performances comparing to machine learning models (Figure 1B-C). To test whether the strategy of featurization caused relatively poor performances, we next evaluated these deep learning models with padding RNA sequences from the 5’ end (Supplementary Figure S2B-C), truncating 5’ end of RNA sequences (Supplementary Figure S2D-E) or using the word2vec method (Supplementary Figure S3A). However, these alternative strategies of featurization didn’t improve performances significantly (Supplementary Figure S2B-C, Supplementary Figure S3C-D), while increased time and memory consumption (Supplementary Figure S3B).

**Table 1.** Evaluation of prediction models for lncRNA subcellular localization by using the lncRNA test set (Test-lncRNA, n = 380)

| Model | Accuracy | Sensitivity | Specificity | MCC | F1 score | AUROC |
| --- | --- | --- | --- | --- | --- | --- |
| RNAlight | 0.72 | 0.76 | 0.68 | 0.45 | 0.74 | 0.78 |
| SVM | 0.69 | 0.77 | 0.60 | 0.37 | 0.72 | 0.75 |
| LR | 0.70 | 0.78 | 0.61 | 0.40 | 0.73 | 0.76 |
| CNN | 0.64 | 0.64 | 0.64 | 0.28 | 0.65 | 0.72 |
| RNN | 0.59 | 0.48 | 0.71 | 0.19 | 0.55 | 0.63 |
| CNN + RNN | 0.65 | 0.65 | 0.65 | 0.31 | 0.66 | 0.71 |

**Table 2.**
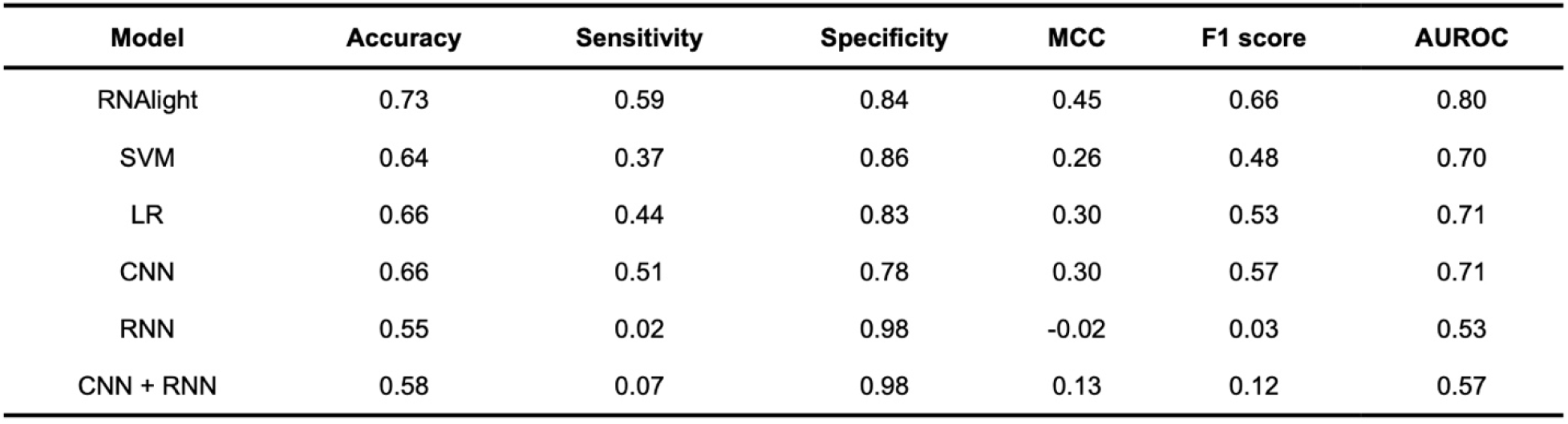
Evaluation of prediction models for mRNA subcellular localization by using the mRNA test set (Test-mRNA, n = 518)

| Model | Accuracy | Sensitivity | Specificity | MCC | F1 score | AUROC |
| --- | --- | --- | --- | --- | --- | --- |
| RNAlight | 0.73 | 0.59 | 0.84 | 0.45 | 0.66 | 0.80 |
| SVM | 0.64 | 0.37 | 0.86 | 0.26 | 0.48 | 0.70 |
| LR | 0.66 | 0.44 | 0.83 | 0.30 | 0.53 | 0.71 |
| CNN | 0.66 | 0.51 | 0.78 | 0.30 | 0.57 | 0.71 |
| RNN | 0.55 | 0.02 | 0.98 | -0.02 | 0.03 | 0.53 |
| CNN + RNN | 0.58 | 0.07 | 0.98 | 0.13 | 0.12 | 0.57 |

To further evaluate the performance of RNAlight, we used test sets (Test-lncRNA, n = 380; Test-mRNA, n = 518) to compare RNAlight with four previously-reported prediction models, including iLoc-lncRNA [22] and lncLocator [21] for lncRNA localization, or iLoc-mRNA [23] and mRNALoc [19] for mRNA localization. As illustrated by the confusion matrix in Figure 2A, RNAlight achieved a more accurate prediction for lncRNA localization than iLoc-lncRNA and lncLocator did, with the highest accuracy, F1 score (Figure 2B) and AUROC (Figure 2C). Similarly, RNAlight also outperformed other compared models in mRNA localization prediction (Figure 2D–2F). In addition, when evaluating with a totally independent dataset of lncRNAs (n = 116, Supplementary figure S4A) and mRNAs (n = 809, Supplementary Figure S4E) from Halo-seq in Hela cell line (Supplementary Table S3, “Methods” section) [31], RNAlight also showed generally better performance than other models on both lncRNA (Supplementary Figure S4B-D) and mRNA (Supplementary Figure S4E-H) subcellular localization prediction.

**Figure 2.**
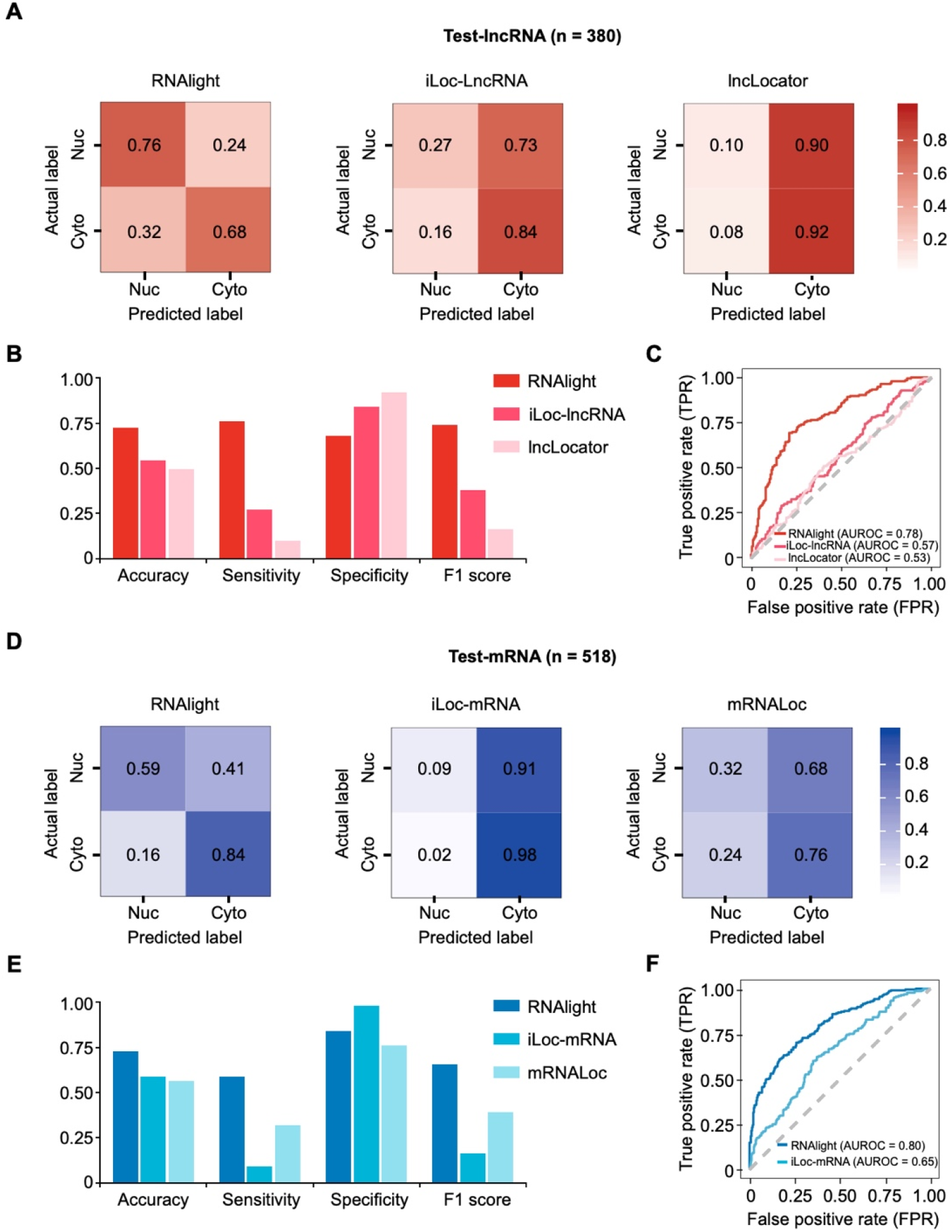
Comparative evaluation of RNAlight and other published models for RNA subcellular localization prediction. The confusion matrix (**A**), accuracy, sensitivity, specificity, F1 score (**B**) and AUROC curve (**C**) of RNAlight, iLoc-LncRNA and lncLocator on predicting lncRNA localization with the lncRNA test set (Test-lncRNA, n = 380). The confusion matrix (**D**), accuracy, sensitivity, specificity and F1 score (**E**) of RNAlight, iLoc-mRNA and mRNALoc on predicting mRNA localization with the mRNA test set (Test-mRNA, n = 518). (**F**) AUROC curves of RNAlight and iLoc-mRNA on predicting mRNA localization with the mRNA test set (Test-mRNA, n = 518).

These results together suggested that the LightGBM-based RNAlight model could accurately predict the different subcellular localizations (nucleus and cytoplasm) for both lncRNAs and mRNAs, which was consistent with the superior performance of LightGBM across a series of benchmark tests in previous studies [24, 32, 33].

### Identifying distinct sequence features for lncRNA or mRNA subcellular localization with Tree SHAP algorithm

As *k*-mers were used as inputs for RNAlight analysis, we next attempted to identify which *k-* mers (sequence features) might play important roles in different (nucleus or cytoplasm) subcellular localizations of lncRNAs and/or mRNAs. In addition to the *k*-mer frequencies (Figure 3A, left), we also used Tree SHAP (SHapley Additive exPlanations) [25] in the LightGBM algorithm for the contribution analysis of *k*-mers in determining RNA nuclear or cytoplasmic localization. In theory, a positive or negative SHAP value suggests a potential role of a given *k*-mer on the nuclear or cytoplasmic localization of a given examined RNA, respectively (“Methods” section). For each *k*-mer, different SHAP values could be quantified to show its distinct contributions on subcellular localization of different examined RNAs (Figure 3A, right). To better access the general effect of each *k*-mer on nuclear or cytoplasmic localization of all examined RNAs, Pearson correlation coefficients (PCCs) between *k*-mer frequencies and SHAP values were calculated (Figure 3A, bottom). In general, a positive PCC represents an overall nuclear localization effect of a given *k*-mer on analyzed lncRNAs or mRNAs, and a negative PCC represents an overall cytoplasmic localization effect of another given *k*-mer on analyzed lncRNAs or mRNAs (“Methods” section). With PCC > 0.5 as cutoff, 399 *k*-mers were determined for nuclear localization of lncRNAs; meanwhile, 501 cytoplasm-related *k*-mers for cytoplasmic localization of lncRNAs were selected by PCC < −0.5 (Figure 3B, and Supplementary Table S4). Importantly, several nuclear localization-related sequence elements of lncRNAs that have been previously identified by a high-throughput screening of short RNA fragments[9, 34] were successfully predicted by RNAlight (Figure 3B). Specifically, a cluster of five-mers, such as CUCCC, CCUCC and ACCUC, were identified with positive PCCs (0.726, 0.727 and 0.537, respectively) by RNAlight (Figure 3B). These five-mers can be tiled across the RCCUCCC motif (where R denotes A/G), which has been previously confirmed associated with lncRNA nuclear localization [9], suggesting the reliable prediction of RNA subcellular localization by RNAlight.

**Figure 3.**
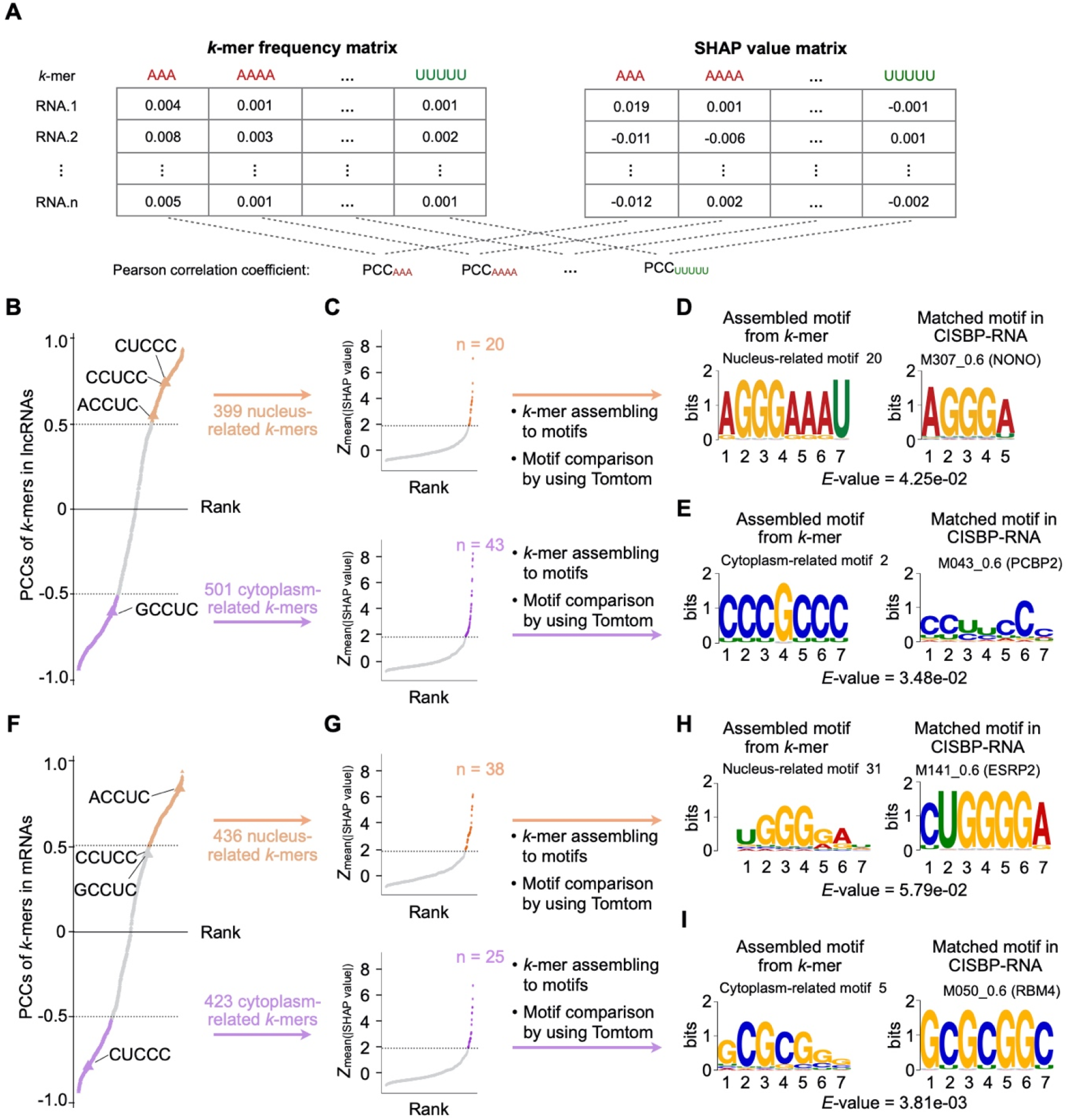
Identification of localization-associated RNA binding proteins for lncRNAs and mRNAs. (**A**) Schematic drawing of PCC (Pearson correlation coefficient) calculation for each given *k*-mer. (**B**) Identification of nucleus- and cytoplasm-related *k*-mers for lncRNAs. The scatter plot shows PCCs of all 1,344 *k*-mers in lncRNAs. (**C**) The scatter plots show *Z*-transformed mean absolute SHAP values of the *k*-mers with PCC > 0.5 (top) or PCC < −0.5 (bottom). (**D-E**) Examples of *k*-mer assembled motifs (left) comparing with known RBP-associated motifs (right) for lncRNAs. (**F**) Identification of nucleus- and cytoplasm-related *k*-mers for mRNAs. The scatter plot shows PCCs of all 1,344 *k*-mers in mRNAs. (**G**) The scatter plots show *Z*-transformed mean absolute SHAP values of the *k*-mers with PCC > 0.5 (top) or PCC < −0.5 (bottom). (**H-I**) Examples of *k*-mer assembled motifs (left) comparing with known RBP-associated motifs (right) for mRNAs.

Given the fact that a spectrum of variable SHAP values could be determined for each particular *k*-mer, we then calculated a *Z*-transformed mean absolute SHAP value for each *k*-mer and used *Z*-score > 1.96 as an additional cutoff to identify most important *k*-mers for RNA subcellular localization among all examined lncRNAs or mRNAs. As shown in Figure 3C and Supplementary Figure S5A and S5B, 20 out of 399 nucleus-related and 43 out of 501 cytoplasm-related *k*-mers were individually identified to play key roles in determining lncRNA nuclear or cytoplasmic localization with *Z*-score > 1.96 (Supplementary Table S4). Of note, *Z*-scores of mean absolute SHAP values of aforementioned five-mers (CUCCC, CCUCC and ACCUC) were < 1.96, possibly due to their limited distribution among a small cluster of lncRNAs[9] but not in thousands of lncRNAs examined in the current study.

It is well known that the interaction of RNA sequences and their associated RBPs is of importance for RNA subcellular localization [9, 35]. We thus aimed to find what types of RBPs could individually bind to these different *k*-mers for distinct RNA subcellular localization. To achieve this goal, we assembled nucleus-related or cytoplasm-related *k*-mers individually to different sequence features groups [26]. Briefly, important localization-related *k*-mers were first mapped back to the RNA sequences. Neighboring *k*-mers were then ligated as candidate sequence features. Consensus sequence features were further obtained by the multiple sequence alignment based on these candidate sequence features (Supplementary Figure S6). From 20 nucleus-related *k*-mers identified by PCC > 0.5 and *Z*-score > 1.96 (Figure 3B), 190 sequence features were obtained by *k*-mer assembling (Supplementary Table S5); from 34 cytoplasm-related *k*-mers identified by PCC < −0.5 and *Z*-score > 1.96 (Figure 3B), eight sequence features were obtained by *k*-mer assembling (Supplementary Table S5). After that, we used Tomtom [36] (“Methods” section) to map these assembled sequence features to known RBP-associated motifs reported in the CISBP-RNA (Catalog of Inferred Sequence Binding Preferences of RNA binding proteins) database [37] (Supplementary Figure S6). As a result, 27 out of 190 nucleus-related sequence features were identified to be associated with 18 RBPs for lncRNA nuclear localization (Supplementary Table S6). For example, NONO (non-POU domain containing octamer binding), a well-studied RBP that preferentially binds RNAs with AGGGA/U elements [38] and participates in paraspeckle formation through binding with nuclear *NEAT1* lncRNA[39], was identified to be associated with lncRNA nuclear localization in this study (Figure 3D). Instead, only two out of eight cytoplasm-related sequence features were identified to be individually associated with two different RBPs for lncRNA cytoplasmic localization (Supplementary Table S6). Between these two RBPs, PCBP2 (poly(rC) binding protein 2), one major cellular poly(rC)-binding protein that is able to bind with C-rich sequences [37] and cooperates with LINC02535 to enhance the stability of mRNA in the cytoplasm [40], was predicted to be involved in regulating lncRNA cytoplasmic localization (Figure 3E).

Similar analyses were parallelly performed for mRNA subcellular localization (Figure 3F–3I). With PCC > 0.5 as cutoff, 436 *k*-mers were determined for nuclear localization of mRNAs, and 423 cytoplasm-related *k*-mers for cytoplasmic localization of mRNAs were selected by PCC < −0.5 (Figure 3F, and Supplementary Table S7). In addition, 38 out of 436 nucleus-related and 25 out of 423 cytoplasm-related *k*-mers were individually identified to play key roles in determining mRNA nuclear or cytoplasmic localization with *Z*-score > 1.96 (Figure 3G, Supplementary Figure S5C-D, and Supplementary Table S7). Of note, no overlap was observed between 20 of important nucleus-related *k*-mers for lncRNAs and 38 of those for mRNAs (Supplementary Figure S5E, left), and only one cytoplasm-related *k*-mer was observed between 43 of important cytoplasm-related *k*-mers for lncRNAs and 25 of those for mRNAs (Supplementary Figure S5E, right). This result (Supplementary Figure S5E) thus suggested distinct *cis*-element features contributing to lncRNA and mRNA subcellular localizations.

In the analysis of identifying what types of RBP could individually bind to different *k*-mers for distinct mRNA subcellular localization, 1,223 sequence features were obtained by *k*-mer assembling (Supplementary Table S8) from 38 nucleus-related *k*-mers identified by PCC > 0.5 and *Z*-score > 1.96 (Figure 3G), and 235 sequence features were obtained by *k*-mer assembling (Supplementary Table S8) from 25 cytoplasm-related *k*-mers identified by PCC < −0.5 and *Z*-score > 1.96 (Figure 3G). After mapping to known RBP-associated motifs by Tomtom, 262 out of 1,223 nucleus-related sequence features were identified to be associated with 54 RBPs for mRNA nuclear localization (Supplementary Table S9). For example, ESRP2 (epithelial splicing regulatory protein 2), an epithelial cell-type specific splicing regulator which was mainly located in nucleus [41] and preferentially binds to RNA with UGGGRAD motif [37], was identified to be linked with the regulation of mRNA nuclear localization (Figure 3H). For the cytoplasmic localization of mRNAs, 67 out of 235 cytoplasm-related sequence features were identified to be associated with 16 RBPs (Supplementary Table S9), including RBM4 (RNA binding motif protein 4) (Figure 3I), an RNA-binding factor involved in mRNA splicing and translation regulation [42] with a tendency to bind with GC-rich sequences [37].

### Applying RNAlight to accurately predict subcellular localizations of various types of RNAs

In contrast to previous models, RNAlight was designed to examine subcellular localization of both mRNAs and lncRNAs. As expected, RNAlight showed a preference of cytoplasmic localization for mRNA transcripts (n = 18,607, GENCODE v30) (the median of Light score = −0.253, Figure 4A), as most mature mRNAs are preferentially transported to the cytoplasm for protein translation [43]. However, a few mRNAs, such as *MLXIPL* and *NLRP6*, were predicted to be the nuclear localization with Light scores of 0.827 and 0.847, respectively (Supplementary Table S10), which are in line with their subcellular localizations in nuclear speckles [8]. Differently, a bimodal distribution with a slightly nuclear preference (the median of Light score = 0.037) of annotated lncRNA transcripts (n = 16,153) was predicted by RNAlight (Figure 4A), suggesting their regulatory roles at multiple components of cells [12], despite a greater proportion of lncRNAs were shown with a nucleus tendency [14, 44]. Accordingly, a set of known nuclear-localized lncRNAs, such as *MALAT1, NEAT1* and *XIST* [39, 45, 46], were predicted by RNAlight with the Light scores of 0.671, 0.899 and 0.854, respectively (Supplementary Table S10). Meanwhile, some known cytoplasmic lncRNAs, such as *ZFAS1* and *SNHG6* [47, 48], were successfully predicted as cytoplasmic localization with Light scores of −0.949 and −0.913 (Supplementary Table S10).

**Figure 4.**
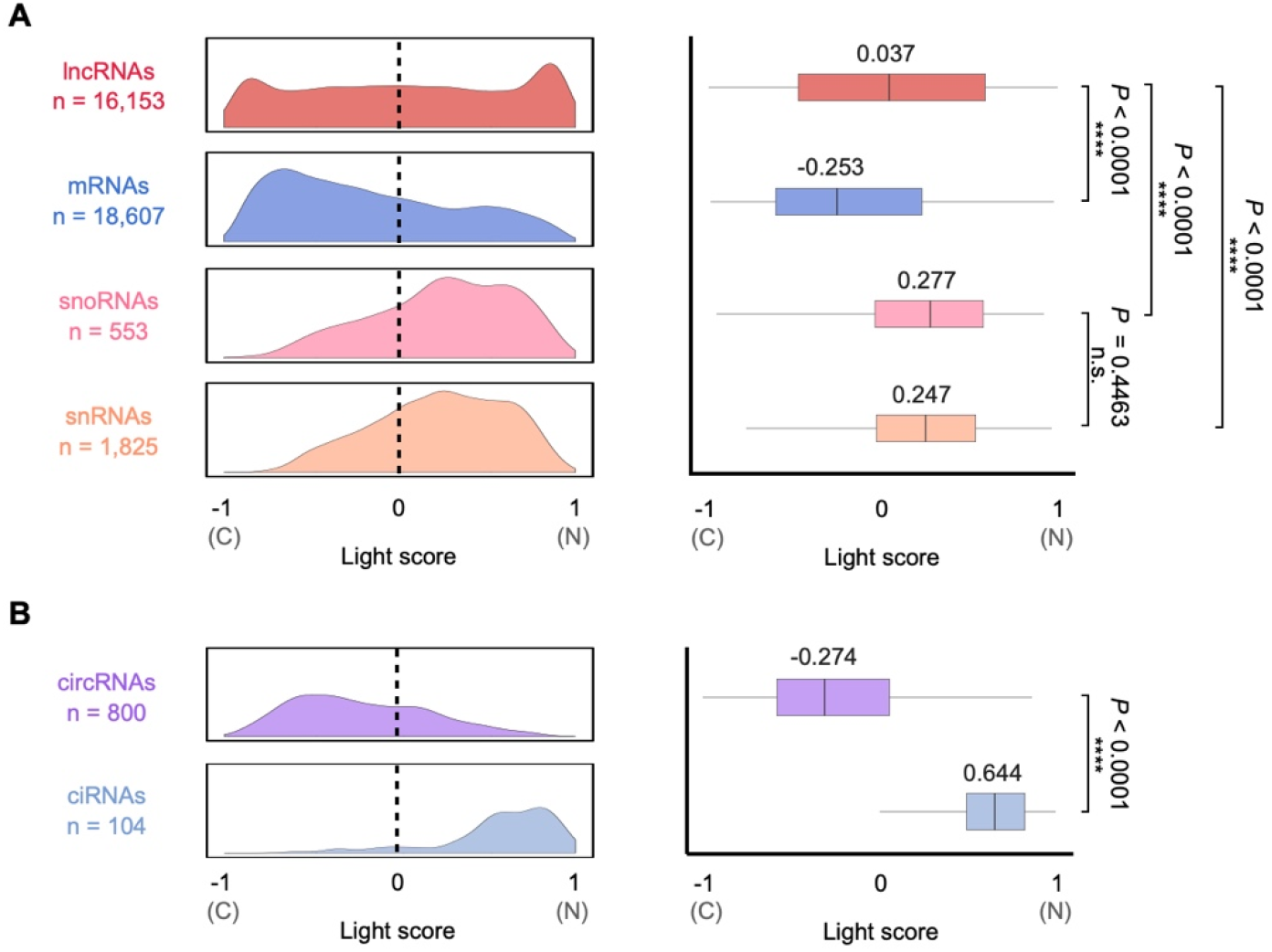
Application of RNAlight to predict subcellular localizations of various types of RNAs. Density (left) and box (right) charts show the distributions of Light scores reported by RNAlight across various types of RNAs, including major transcripts of lncRNA, mRNA, snoRNA and snRNA from GENCODE v30 annotation (**A**), and circRNAs and ciRNAs identified in PA1 cells from Zhang et al [61] (**B**). The range of Light score is from −1 to 1, wherein the interval of −1 to 0 or 0 to 1 indicates cytoplasmic or nuclear localization of RNA, respectively. Statistical testing was performed with two-sided Welch’s *t*-test; n.s., not significant.

To further evaluate the generality of RNAlight for the subcellular localization prediction on different types of RNAs, we extended the analysis to other RNA species that were not used for the model training, including small nucleolar RNAs (snoRNAs), small nuclear RNAs (snRNAs), and circular RNAs. It is well known that snoRNAs are distributed in eukaryotic nucleolus for rRNA/tRNA methylation or other RNA modification [49], and snRNAs are localized in the nucleoli and nucleolus as main components of spliceosome [50]. Correspondingly, RNAlight accurately predicted their nuclear localization of snoRNAs (n = 553, the median of Light score = 0.277) and snRNAs (n = 1,825, the median of Light score = 0.247) (Figure 4A). In addition, two major types of spliceosome-dependent circular RNAs, circRNAs from back-spliced exons and circular intronic RNAs (ciRNAs) from spliced intron lariats, were recently rediscovered at a genome-wide level in eukaryotes but with different subcellular localization [51, 52]. It has been reported that circRNAs were generally localized in cytoplasm [53], involving in innate immunity [54, 55], cell proliferation [56] and neuronal function [57], while ciRNAs were preferentially retained in nucleus to regulate Pol II transcription [58, 59]. By using highly expressed circRNAs (FPB > 0.5, n = 800) and ciRNAs (FPB > 0.2, n =104) identified by the CLEAR pipeline [60] in PA1 cells from published ribo– RNA-seq (GEO: GSE73325) [61] (“Methods” section) as inputs, we set up to examine whether RNAlight could be further extended for circular RNA subcellular localization prediction. As shown in Figure 4B, RNAlight successfully predicted the cytoplasmic localization of circRNAs (the median of Light score = −0.274) and the nuclear localization of ciRNA (the median of Light score = 0.644). In contrast, other RNA subcellular localization prediction tools, such as iLoc-lncRNA, lncLocator, iLoc-mRNA and mRNALoc, failed to accurately reveal the subcellular localization of cytoplasmic circRNAs and nuclear ciRNAs (Supplementary Figure S7). These results together indicated RNAlight as a reliable and universal model for the subcellular localization prediction of distinct RNA species.

## Discussion

Here, we report RNAlight, a machine learning model based on LightGBM for precise RNA localization prediction (Figure 1 and Figure 2). In our hands, all examined machine learning models (RNAlight, SVM and Logistic Regression) outperformed deep learning models (CNN, RNN, and CNN+RNN) in the prediction of RNA subcellular localization. Applying different strategies of featurization for deep learning models did not significantly improve their performances (Supplementary Figure S2–S3). The poor performance by deep learning models is possibly due to the relatively small scale of the dataset for model development. In the future, we assume that better deep learning models can be developed with fast-accumulated large-scale datasets. Nevertheless, by integrating the Tree SHAP algorithm and *k*-mer assembly into the RNAlight model, determinant sequence features and their associated RBPs were identified to contribute to distinct mRNA or lncRNA subcellular localizations (Figure 3). Importantly, RNAlight was also shown to be a reliable model for the subcellular localization prediction of various RNAs, including mRNAs, lncRNAs, snoRNAs, snRNAs, circRNAs and ciRNAs (Figure 4), suggesting its broad application for the localization prediction of various types of RNAs.

Compared to some other methods [19–23], RNAlight has two characteristics for the prediction of RNA subcellular localization. On the one hand, by integrating the Tree SHAP algorithm and *k*-mer assembly, RNAlight not only predicts RNA subcellular localization, but also effectively identifies distinct sequence features for their different subcellular (nuclear or cytoplasmic) localization. Similar to previous studies showing that certain *cis*-elements in the 5’ end or 3’ end of RNAs contribute to their subcellular localization [7, 62–64], many *k*-mers identified by RNAlight in this study were also shown to be enriched in 5’ or 3’ ends of their host RNAs (Supplementary Figures S8–S11), indicating that RNAlight could indeed learn important sequence features that determine RNA subcellular localization. Further analyses showed that these sequence features could be associated with distinct RBPs to affect RNA subcellular localization (Figure 3). On the other hand, RNAlight has been also extended to predict subcellular localization of various RNA species, with reliable results that are consistent with previous observations from experimental examination (Figure 4), suggesting the robustness of RNAlight for the prediction of RNA subcellular localization. Interestingly, RNAlight could successfully differentiate RNA circles with exon or intron origins in their distinct subcellular localization. Different to cytoplasmic localization of most circRNAs from back-spliced exons, ciRNAs that are produced from spliced intron lariats were predicted to be preferentially located in nucleus (Figure 4B), consistent with experimental lines of evidence [58, 59]. Similarly, RNAlight showed that the inclusion of intron sequences in lncRNA, mRNA and circRNA could substantially change their subcellular localization from cytoplasm to nucleus (Supplementary Figure S12), in line with the previous study that RNA transcripts with retained introns are considered as incompletely spliced forms and generally retained in nucleus [65].

Despite of these advantages, RNAlight might also simplify the prediction of RNA subcellular localization by only inputting features from RNA primary sequences. However, when adding additional features of predicted RNA secondary structure and the compositional information, the performances of both machine learning and deep learning models were not significantly improved (Supplementary Figure S13, “Methods” section). Since both RNA secondary structure and compositional information were predicted or obtained from RNA primary sequences, we speculated that sequence features might be sufficient to achieve RNA subcellular localization prediction in this scenario. In the future, experimental lines of evidence on RNA secondary structure and modification could be further considered to improve this model.

In addition, given that our study aims to identify general sequence features associated with RNA subcellular localization, RNAlight was trained with RNAs only showing single subcellular localization as many other prediction models did [19, 21–23]. It is thus a common limitation of most existing computational tools for RNA subcellular localization prediction. Nevertheless, we did not rule out the possiblitity that RNAs with distinct subcellular localizations across different cell types or states are functionally important. Identification of other factors, such as associated partners (RBPs, U1 snRNA, etc) and/or cellular contexts in various conditions, is warranted to understand how distinct subcellular localizations could be regulated for a given RNA. Finally, RNAlight only focused on predicting two major RNA subcellular localizations, nucleus and cytoplasm, due to the limited datasets. We expected that detailed RNA subcellular localization could be dissected with more datasets containing high-resolution RNA localization information, such as ER, mitochondria or nucleolus.

Taken together, we reported the RNAlight model based on LightGBM to precisely predict nuclear or cytoplasmic localization of mRNAs and lncRNAs, and further identified important *k*-mer features and RBPs possibly involved in their subcellular localizations. In the future, additional datasets with extra features, including but not limited to RNA secondary structure and/or chemical modification, can be included to train a better model for characterizing complex and dynamic RNA subcellular localization.

## Materials and methods

### Constructing a combined library of lncRNAs and mRNAs labeled with distinct nuclear or cytoplasmic localizations

To generate a universal model for both lncRNA and mRNA subcellular localization prediction, we first collected reported datasets with lncRNA and/or mRNA subcellular localization information, and further combined them together for a combined library of lncRNA and mRNA subcellular localization.

On the one hand, several lncRNA subcellular localization datasets, including LncATLAS [17], CeFra-seq [15] and APEX-Seq [16], were collected for this study (Supplementary Figure S1A). Due to distinct methodologies in these datasets for subcellular localization analysis, different filtering strategies were then implemented in this study to select nuclear or cytoplasmic lncRNAs: 1) LncATLAS database records localization information of 6,768 lncRNAs across 14 cell lines[17]. Here, the mean value of cytoplasmic/nuclear concentration index (*CN–RCI_mean_*) across 13 cell lines (excluding H1 cell due to its low correlation to other cell lines, data not shown) was used for filtering: lncRNAs with (*CN-RCI_mean_*) < −2 (n = 1,857) were considered as nuclear localization, while those with (*CN–RCI_mean_*) > 0 (n = 1,440) were considered as cytoplasmic localization. 2) CeFra-seq extracts cell fractions of cytosol, insoluble, membrane and nucleus for high-throughput sequencing and RNAs localized in these diverse cell fractions could be identified, and 14,746 lncRNAs were detected at these four cell fractions in HepG2 cells [15]. From the CeFra-seq dataset, 1,621 highly-expressed lncRNAs with fragments per kilobase per mapped fragments (FPKM) ≥ 1 in at least one cellular fraction were obtained for subsequent analysis. Accordingly, *CR* (cytoplasmic ratio) was used to distinguish nuclear and cytoplasmic lncRNAs in the CeFra-seq dataset, computed as below:

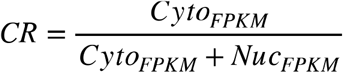

Here, *Nuc_FPKM_* is the FPKM value of a lncRNA in the nuclear fraction and *Cyto_FPKM_* is the maximum FPKM value of the lncRNA in the cytosol, insolubles or membrane fraction. With these criteria, 435 lncRNAs with *CR* < 0.4 were considered with the preference of nuclear localization, and 844 lncRNAs with *CR* > 0.6 were considered with the preference of cytoplasmic localization. 3) APEX-Seq provides a practical methodology to identify RNA in distinct subcellular locales, based on APEX2-mediated proximity biotinylation of endogenous RNAs in the presence of biotin-phenol (BP) and H_2_O_2_ and following poly(A)+ RNA sequencing [16]. RNA localized in one subcellular locale could be identified by calculating the fold change of the H_2_O_2_-treated sample to the untreated control sample in HEK293T cells. Here, among all lncRNAs identified in the APEX-Seq dataset (n = 61), those with log2(fold change) ≥ 0.75 in at least one component of the nucleus, nucleolus, lamina and nuclear pore were considered as nuclear lncRNAs (n = 42) and those with log2(fold change) ≥ 0.75 in at least one component of the cytoplasm, ER (endoplasmic reticulum) membrane, ER lumen and outer mitochondrial membrane were considered as cytoplasmic lncRNAs (n = 5).

After the aforementioned filtering, lncRNAs from these three resources were combined and lncRNAs with inconsistent but multiple localizations were removed to generate a combined library with 1,986 nuclear and 1,806 cytoplasmic (totally 3,792) lncRNAs.

On the other hand, mRNAs with different subcellular localization were collected from CeFra-seq and APEX-Seq datasets (Supplementary Figure S1B). Similar processing parameters for CeFra-seq and APEX-Seq data were applied to select mRNAs with different subcellular localization. From the CeFra-seq dataset, 1,789 and 2,040 mRNAs were selected with the preference of nuclear or cytoplasmic localization, respectively. From the APEX-Seq dataset, 1,145 and 1,261 mRNAs were selected with the preference of nuclear or cytoplasmic localization. Finally, mRNAs with different subcellular localization from these two resources were combined and those with inconsistent but multiple localizations were filtered out, leading to a total of 5,180 mRNAs with nuclear (2,256) or cytoplasmic (2,924) labels.

Collectively, by stringent filtering, we constructed a combined library of lncRNAs (n = 3,792) and mRNAs (n = 5,180) labeled with distinct nuclear or cytoplasmic localizations. These RNAs were randomly split into training sets (Training-lncRNA, n = 3,412; Training-mRNA, n = 4,662) and test sets (Test-lncRNA, n = 380; Test-mRNA, n = 518) with a 9:1 ratio for model training and evaluation.

### Constructing an independent dataset of lncRNAs and mRNAs labeled with distinct nuclear or cytoplasmic localizations from Halo-seq

A totally independent dataset containing lncRNA and mRNA subcellular localization information from Halo-seq in HeLa cell line [31] was collected to further evaluate RNAlight with other published models. With adjusted *P*-value < 0.05 and absolute log_2_(fold change) ? 0.5 as cutoff, H2B-Halo enriched- and Halo-p65 depleted-RNAs were considered as nuclear localization; while H2B-Halo depleted- and Halo-p65 enriched-RNAs were considered as cytoplasmic localization. After removing redundant and bi-localized ones, 116 lncRNAs and 809 mRNAs with the nuclear or cytoplasmic label were individually obtained (Supplementary Table S3) for model evaluation.

### Transcript selection

The files recording the information of RNA localization and Ensembl gene IDs of their parental genes were directly downloaded from public resources [15–17, 31]. For each RNA, we selected the major splice annotation (the −001 isoform) with the GENCODE v30 annotation file as the previously reported method [66] to obtain its primary sequence for subsequent analyses.

### Featurization of RNA primary sequences

The strategy of featurization for machine-learning based models was similar to that in the previous publication [26]. Briefly, frequencies of 1,344 *k*-mers (*k* equals to 3, 4, 5) that permute four nucleotides (A, T, G, C) were firstly computed to show their presence in each specific RNA. These frequencies were further normalized by the RNA length and integrated as a *k*-mer frequency matrix, which can characterize each RNA with 1,344 distinct features.

For featurization in deep-learning based models, each RNA sequence was processed to a fixed length (lncRNA, 4,000nt; mRNA, 9,000nt). Specifically, lncRNAs shorter than 4,000nt or mRNAs shorter than 9,000 nt in length were padded with ‘N’ at their 3’ ends, which covered 95% of lncRNAs and mRNAs according to the 95 percentile of the RNA length (lncRNA, 3,825nt; mRNA, 8,469nt). The rest of lncRNAs and mRNAs with longer lengths were truncated to the same length (lncRNA, 4,000nt; mRNA, 9,000nt). After that, all RNAs with the fixed length were converted to tensor by one-hot encoding as input, where each nucleotide was transformed to a binary vector: A (1, 0, 0, 0), T (0, 1, 0,0), G(0, 0, 1, 0), C(0, 0, 0, 1), N(0, 0, 0, 0).

### Featurization of RNA predicted secondary structure and compositional information

Secondary structures of mRNAs and lncRNAs were predicted by Vienna RNAfold [67] with default parameters using RNA primary sequences. The bpRNA algorithm [68] was then used to annotate predicted structures into different types, including stem (S), hairpin loop (H), multi-loop (M), internal loop (I), bulge (B), external loop (X) and end (E). The secondary structure matrix was constructed to contain the lengths and ratios of different structure types and the minimum free energy (MFE) of examined RNAs (Supplementary Figure S13A, middle panel).

For compositional information of RNA, the GC content, AUGC ratio, GC skew and Z-curve of RNA sequence were calculated by following mathematical formulas:

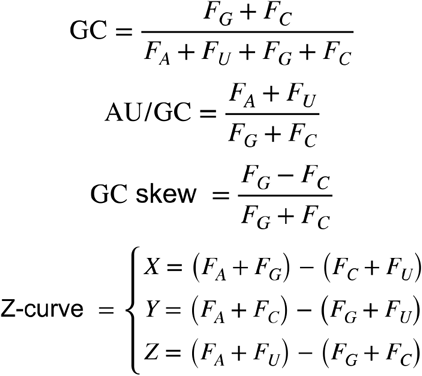

Where F_x_ represents the frequency of each nucleotide (A, U, G, C).

The *k*-mer frequency matrix (representing sequence feature, Supplementary Figure S13A, left panel), the secondary structure matrix (representing structure feature, Supplementary Figure S13A, middle panel) and the composition matrix (representing composition feature, Supplementary Figure S13A, right panel), have been combined as the new input to train the RNAlight model (Supplementary Figure S13A). For comparison, the structure and composition features were also added to train the CNN model with a shared dense layer (Supplementary Figure S13B).

### Construction of RNAlight with LightGBM framework

LightGBM is a new machine-learning implementation of gradient enhanced decision tree (GBDT) with gradient-based one-side sampling and exclusive feature building, which has several advantages, including faster training speed, higher efficiency and lower memory usage. Here, we used the LightGBM Python package (version 3.1.1.99) to train the RNAlight model by inputting the *k*-mer frequency matrix and labeled subcellular localizations from training sets (Training-lncRNA, n = 3,412; Training-mRNA, n = 4,662).

Five-fold cross-validation based on RandomizedSearchCV was performed to select the optimal hyperparameters for LightGBM. We searched 1,000 combinations chosen from following hyperparameter configurations: learning rate (chosen from [0.1, 0.05, 0.02, 0.01]), the number of estimators (chosen from 24 points that were evenly spaced between 100 and 2,400), the maximum depth of the individual estimators (chosen from [2, 3, 4, 5, 10, 20, 40, 50]), the minimum number of data in one leaf (chosen from 22 points that were evenly spaced between 1 and 44), the fraction of subset on each estimator (chosen from [0.2, 0.3, 0.4, 0.5, 0.6, 0.7, 0.8, 0.9]), the frequency of bagging (chosen from [0, 1, 2]), the penalty of L1 regularization (chosen from [0, 0.001, 0.005, 0.01, 0.1]) and the penalty of L2 regularization (chosen from [0, 0.001, 0.005, 0.01, 0.1]).

### Construction of support vector machine (SVM) model and logistic regression model

SVM and logistic regression models were both performed by Python scikit-learn package (version 0.20.3). For SVM, we searched 60 combinations through the following hyperparameter configurations: the kernel type (chosen from [“linear”, “rbf’]), the penalty parameter *C* (chosen from [0.01, 0.1, 1, 10, 100]) and the kernel coefficient *γ* (chosen from [0.001, 0.005, 0.1, 0.5,1, 2]). For logistic regression, the penalty of L2 regularization was chosen from [1e-3, 5e-3, 1e-2, 0.05, 0.1, 0.5, 1, 5, 10, 50, 100, 500, 1000] to optimize hyperparameters. Five-fold crossvalidation based on RandomizedSearchCV was also utilized to select the appropriate set of hyperparameters.

### Construction of deep learning-based models

CNN (convolutional neural network), RNN (recurrent neural network) and a combinatorial model of CNN and RNN (CNN+RNN) were applied to train deep learning-based models under the Tensorflow (version 2.0.0) backend in Python (version 3.6.12).

CNN was developed using two convolutional layers and one dense layer with the following hyperparameters: the number of filters [32, 64, 128] in the first convolutional layer and the number of units [256, 512, 1,024] in the dense layer. RNN was developed using one Bidirectional LSTM layer and one dense layer with flowing hyperparameters: the number of hidden units [16, 32, 64] in Bidirectional LSTM layer and the number of units [256, 512, 1,024] in the dense layer; the combinatorial model of CNN and RNN (CNN+RNN) was developed using two convolutional layers, one Bidirectional LSTM layer and one dense layer with following hyperparameters: the number of filters [32, 64, 128] in first convolutional layer and the number of hidden units [16, 32, 64] in Bidirectional LSTM layer. We chose the model showed the highest mean AUROC (area under the receiver operating characteristic curve) in the five-fold cross-validation.

### Evaluation of model performance

We used test sets, including Test-lncRNA (n = 380) and Test-mRNA (n = 518), which were excluded during model training, to evaluate model performances on predicting subcellular localizations of lncRNA and mRNA, respectively. Based on the confusion matrix from actual and predicted labels, models were assessed with the following indicators:

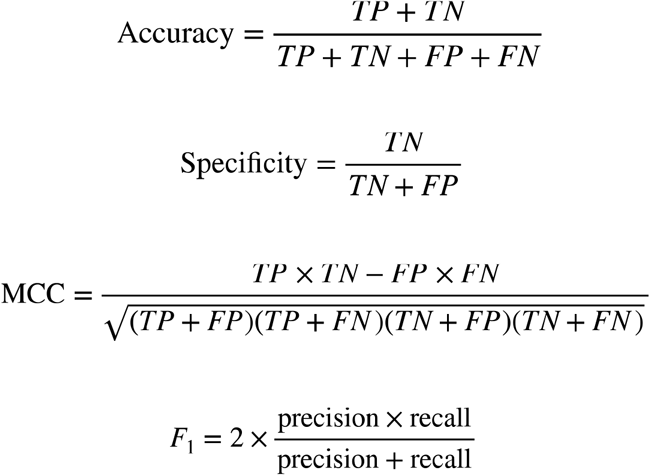

The AUROC was also calculated among the models except for mRNALoc because of its unsuitable possibilities from the output.

In our hands, the LightGBM-based RNAlight model developed in this study showed the best performance for both lncRNA and mRNA subcellular localization prediction.

### Identification of nucleus- and cytoplasm-related *k*-mers

Tree SHAP [25] (SHapley Additive exPlanations) was used to measure *k*-mer contribution to RNA subcellular localization. In the Tree SHAP method, the SHAP value, which is calculated on the basis of a game theoretic Shapley value for optimal credit allocations, is assigned to each identified *k*-mer in a specific RNA from RNAlight.

Nucleus- and cytoplasm-related *k*-mers were identified with two criteria: 1) For each given *k*-mer, Pearson correlation coefficients (PCCs) between the *k*-mer frequencies and SHAP values of each given *k*-mer in all examined RNAs are obtained to reflect its impact on the model output. 2) The *Z*-score of mean absolute SHAP value is used to provide a general overview of its importance on the subcellular localization of all examined RNAs. Taken together, *k*-mers with PCC > 0.5 and *Z*-score > 1.96 suggest to be associated with nuclear localization, and ones with PCC < −0.5 and *Z*-score > 1.96 suggest to be associated with cytoplasmic localization.

### Identification of known RBP-associated motifs related to RNA subcellular localization

Nucleus- and cytoplasm-related *k*-mers were separately assembled to consensus sequence features by the previously reported method [26]. Briefly, we tiled these *k*-mers back to each transcript (e.g., nucleus-related *k*-mers were tiled to the nucleus-localized transcripts) and joined consecutive *k*-mers together to form longer sequences as candidate sequence features. These candidate sequence features were merged to identify consensus sequences by multiple sequence alignment. Consensus sequence features were then mapped to known human RBP position weight matrices (PWMs) in CISBP-RNA database [37], which consists of RNA motifs and specificities to RBPs, by Tomtom [36] (version 5.3.0, https://meme-suite.org/meme/meme_5.3.0/tools/tomtom) to identify known RBP-associated motifs related to RNA subcellular localization.

### Calculation of Light score

To use RNAlight for the prediction of a given RNA, we scaled the output of the RNAlight model as Light score:

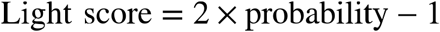

Here, the probability is the original output from RNAlight ranging from 0 to 1, representing the probability of nuclear localization about the input RNA. Scaled Light scores range from −1 to 1, of which a given Light score in the interval (−1, 0) indicates cytoplasmic localization or in the interval (0, 1) indicates nuclear localization.

### Selection of circular RNAs

Highly expressed circRNAs without retained introns (FPB > 0.5, n = 800) and ciRNAs (FPB > 0.2, n =104) in PA1 cells from published ribo– RNA-seq (GEO: GSE73325) [61] were identified by CLEAR pipeline [60] in this study, and these circular RNAs were further used to evaluate the performance of RNAlight. Circular RNA sequences were stretched by Bedtools (version 2.28.0).

## Acknowledgement

We thank Yang laboratory for discussion, and previous lab members, Zheng Luo, He-Na Zhang and Meng-Ran Wang for their early tests on this project.

## Data availability

All scripts used in this project are currently available at https://github.com/YangLab/RNAlight, including RNAlight model and related codes.

## Supplementary figures and figure legends

**Supplementary Figure S1.**
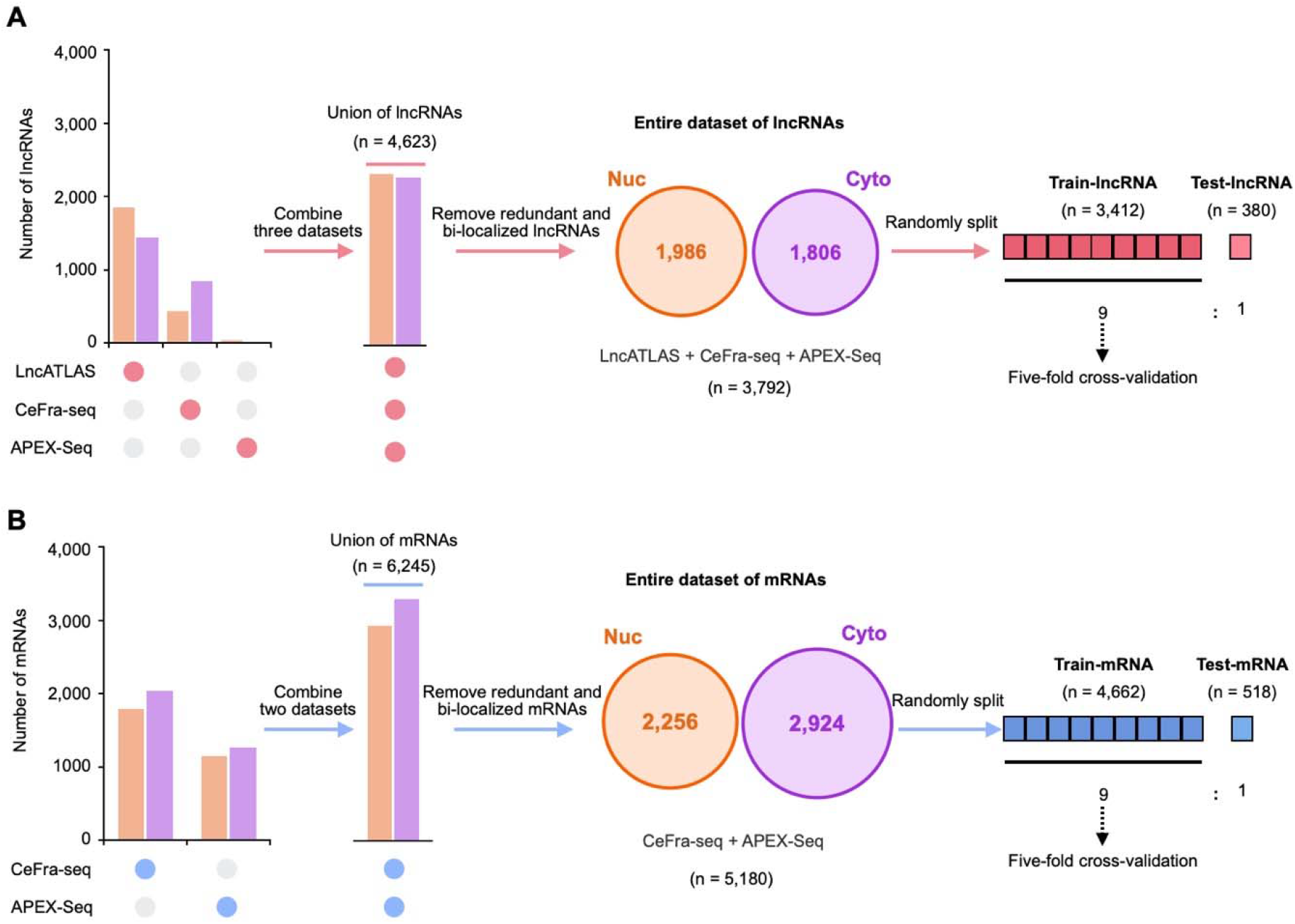
A combined library of RNAs labeled with subcellular localization. Nucleus (orange) and cytoplasm (violet) localized RNAs are extracted from multiple datasets, including LncATLAS, CeFra-seq and APEX-Seq. After removing redundant and multi-localized ones, a combined collection of lncRNAs (**A**, n = 3,792) and mRNAs (**B**, n = 5,180) labeled with distinct nuclear or cytoplasmic localization was obtained, and individually split into training sets and test sets with a 9:1 ratio for model training and evaluation, respectively.

**Supplementary Figure S2.**
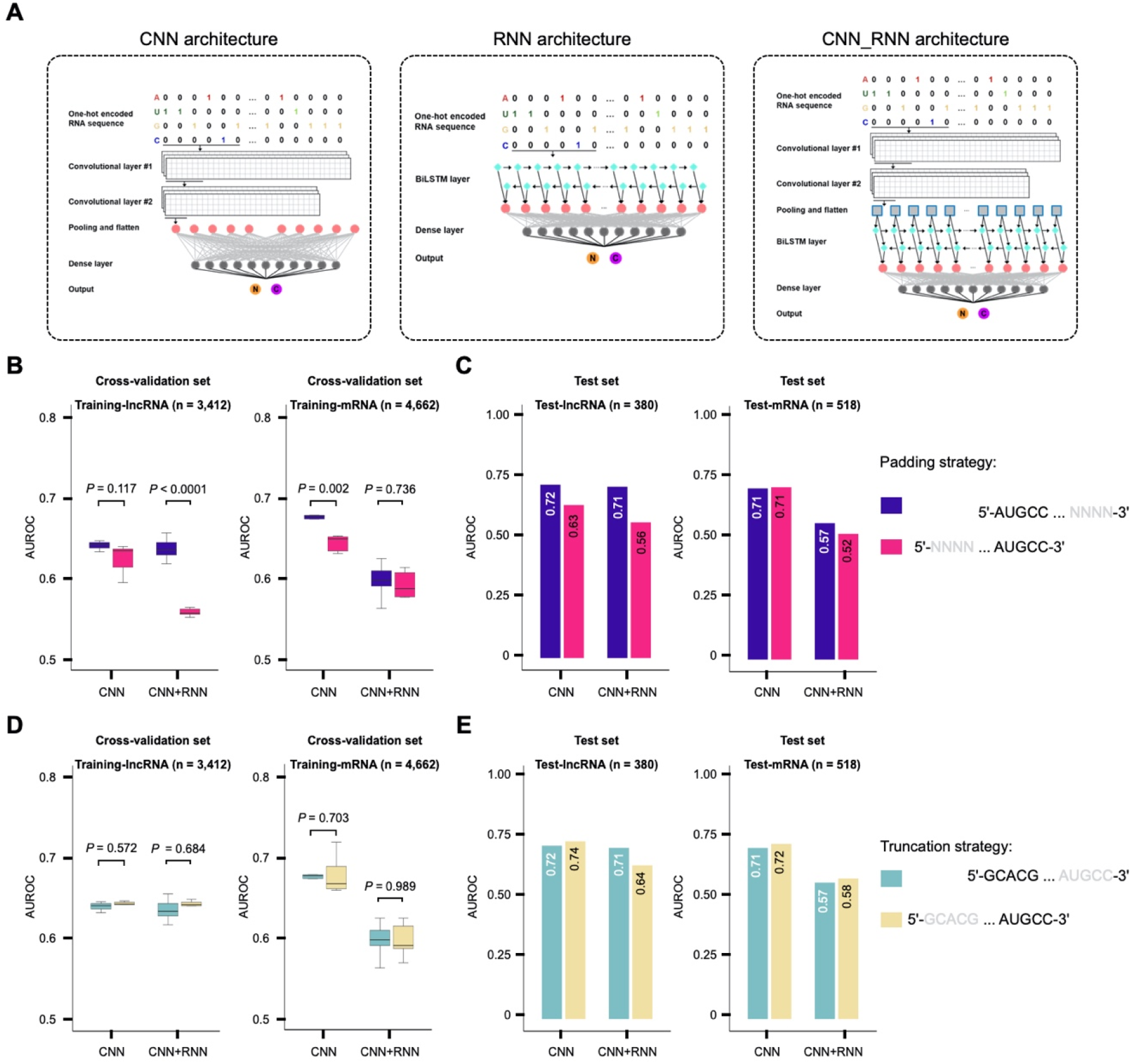
Comparative evaluation of different padding and truncating strategies in deep learning models for RNA subcellular prediction. (**A**) Diagrams of three deep learning models used in this study. (**B**) Cross-validation of models with different padding strategies for lncRNA (left) and mRNA (right) subcellular localization. Each dot represents the area under the receiver operating characteristic curve (AUROC) from five-fold cross-validation (total AUROC, n = 5). Statistical testing was performed with one-sided Welch’s *t*-test. In the box plots, the 25th, 50th and 75th percentiles are indicated as the top, middle, and bottom lines, respectively; whiskers represent the 10th and 90th percentiles, respectively. (**C**) Evaluation of models with different padding strategies for lncRNA (left) and mRNA (right) subcellular localization by using the test sets. Note: RNN model didn’t pad RNA sequence because it can handle input with variable lengths. (**D**) Cross-validation of models with different truncating strategies for lncRNA (left) and mRNA (right) subcellular localization. (**E**) Evaluation of models with different truncating strategies for lncRNA (left) and mRNA (right) subcellular localization by using the test sets.

**Supplementary Figure S3.**
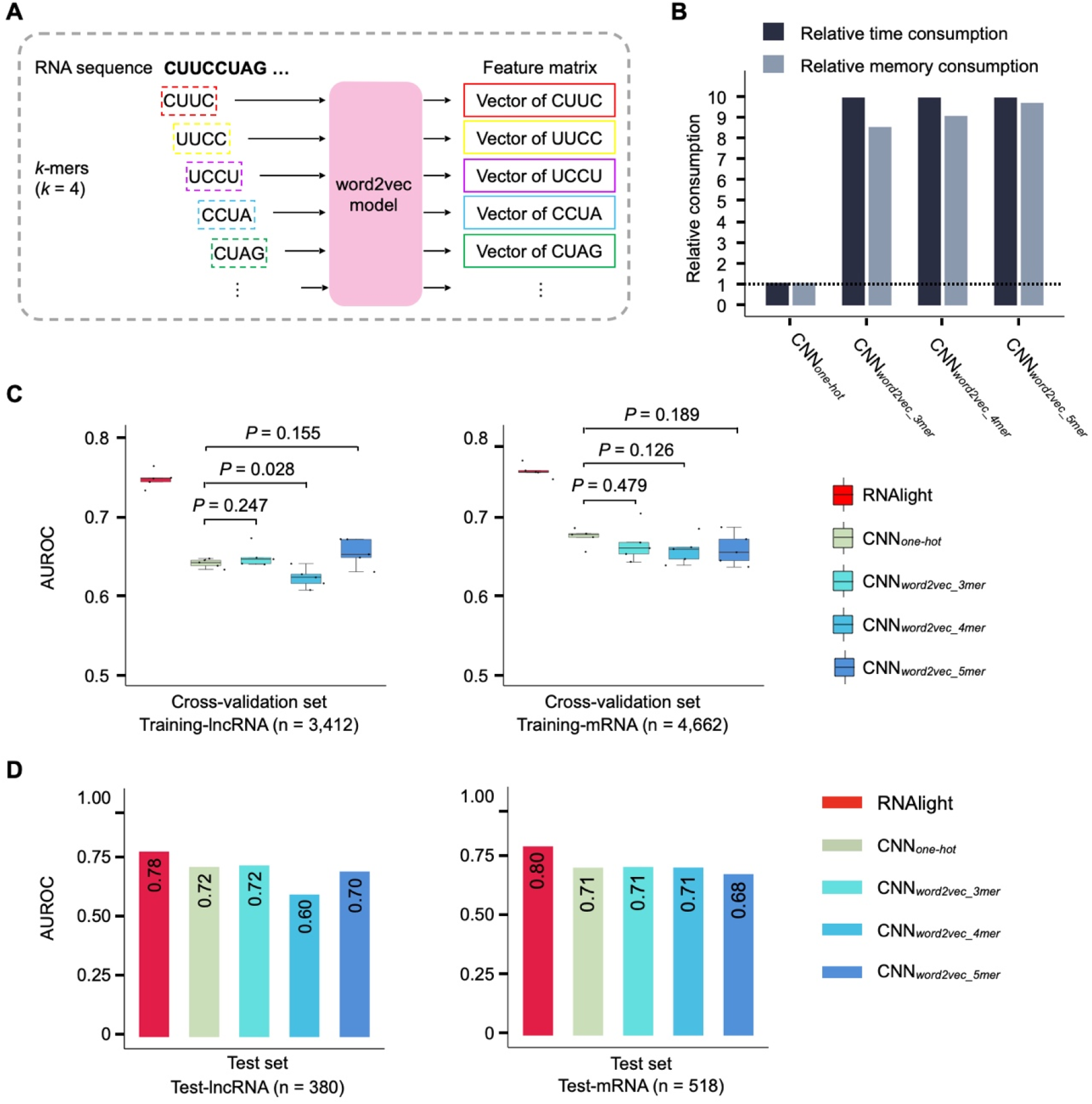
Encoding RNA sequences with word2vector method. (**A**) Schematic drawing of encoding RNA sequences with word2vector. (**B**) Comparison of time and memory consumption between one-hot and word2vector methods in CNN framework. (**C**) Cross-validation of models with different encoding strategies for lncRNA (left) and mRNA (right) subcellular localization. Each dot represents the area under the receiver operating characteristic curve (AUROC) from five-fold cross-validation (total AUROC, n = 5). Statistical testing was performed with one-sided Welch’s *t*-test. In the box plots, the 25th, 50th and 75th percentiles are indicated as the top, middle, and bottom lines, respectively; whiskers represent the 10th and 90th percentiles, respectively. (**D**) Evaluation of models with different encoding strategies for lncRNA (left) and mRNA (right) subcellular localization by using the test sets.

**Supplementary Figure S4.**
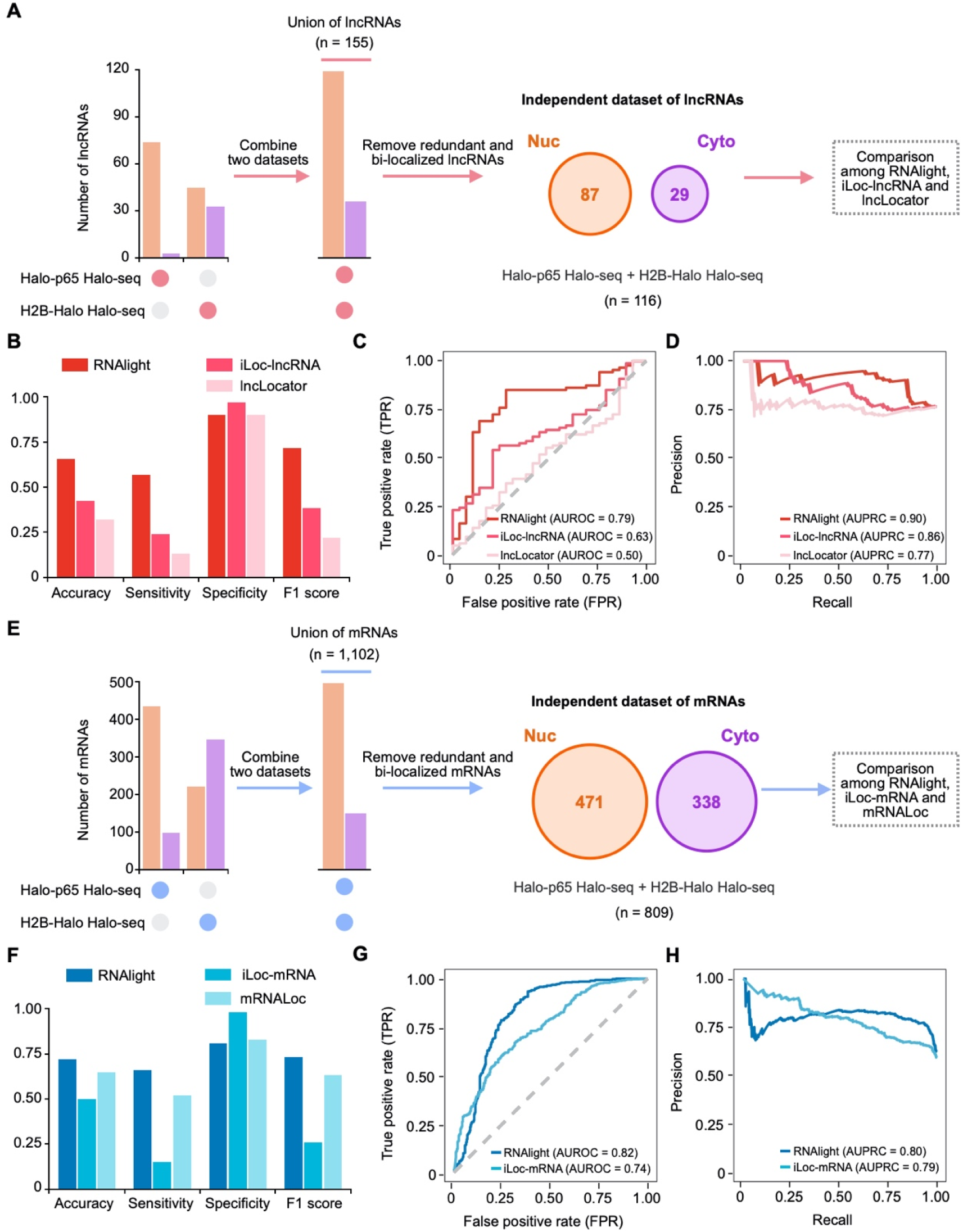
Comparison of RNAlight and published models with an independent dataset. A totally independent dataset from Halo-seq was used for lncRNA (A-D) or mRNA (E-H) subcellular localization prediction by RNAlight and some published models.

**Supplementary Figure S5.**
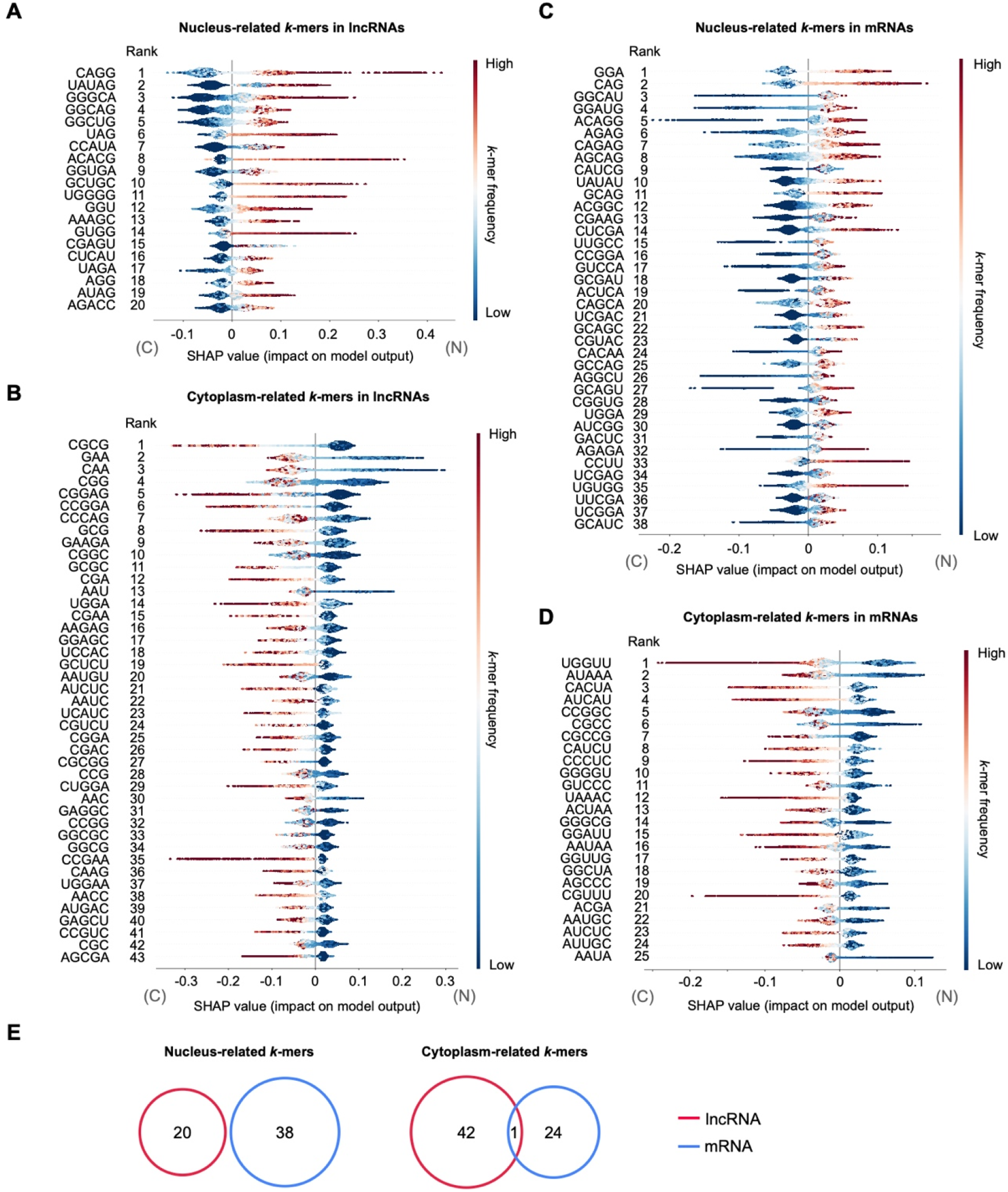
Identification of important localization-associated *k*-mers for lncRNAs. Violin plots show SHAP values of nucleus-related *k*-mers (n = 23) (**A**) and cytoplasm-related *k*-mers (n = 43) (**B**) in lncRNAs, or SHAP values of nucleus-related *k*-mers (n = 38) (**C**) and cytoplasm-related *k*-mers (n = 25) (**D**) in mRNAs. Each dot corresponds to one SHAP value of a given *k*-mer in a given RNA. The color of a dot represents the frequency of a given *k*-mer in each RNA. Positive and negative SHAP values of *k*-mers indicate their contribution to nuclear or cytoplasmic subcellular localization, respectively. (**E**) Overlap of localization-related *k*-mers between lncRNAs and mRNAs.

**Supplementary Figure S6.**
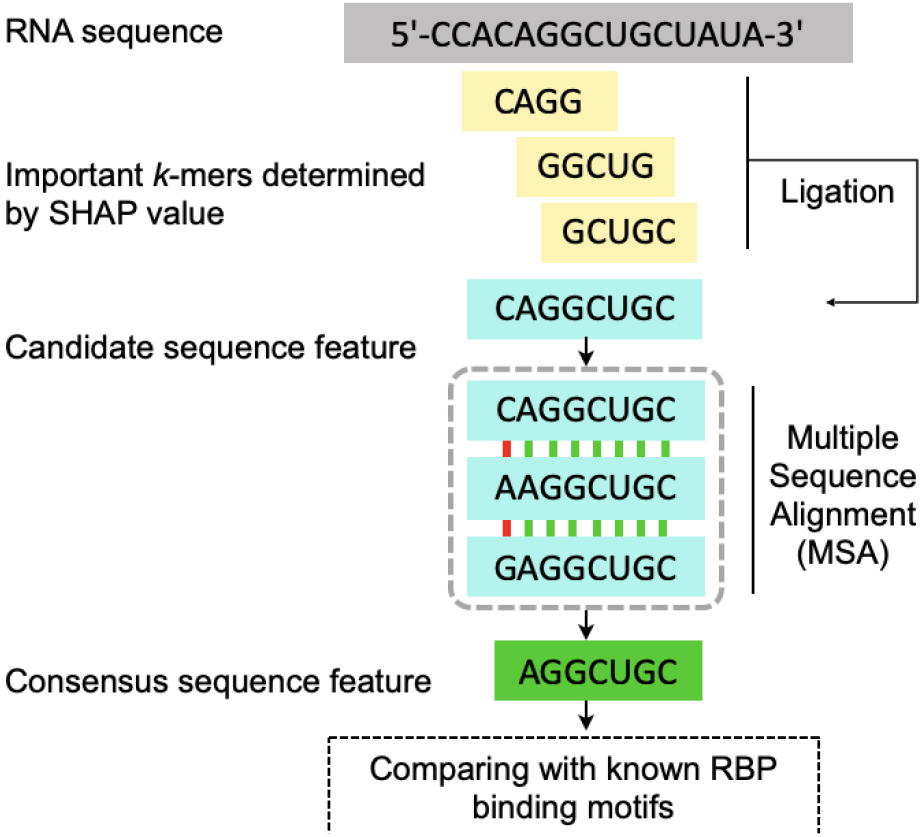
Schematic drawing of assembling *k*-mers to the consensus sequence features. Important *k*-mers were tiled back to each transcript and joined together to form longer sequences as candidate sequence features, which were merged to identify consensus sequences by multiple sequence alignment. Consensus sequence features were further mapped to known human RBP position weight matrices (PWMs) in CISBP-RNA database to identify known RBP-associated motifs related to RNA subcellular localization.

**Supplementary Figure S7.**
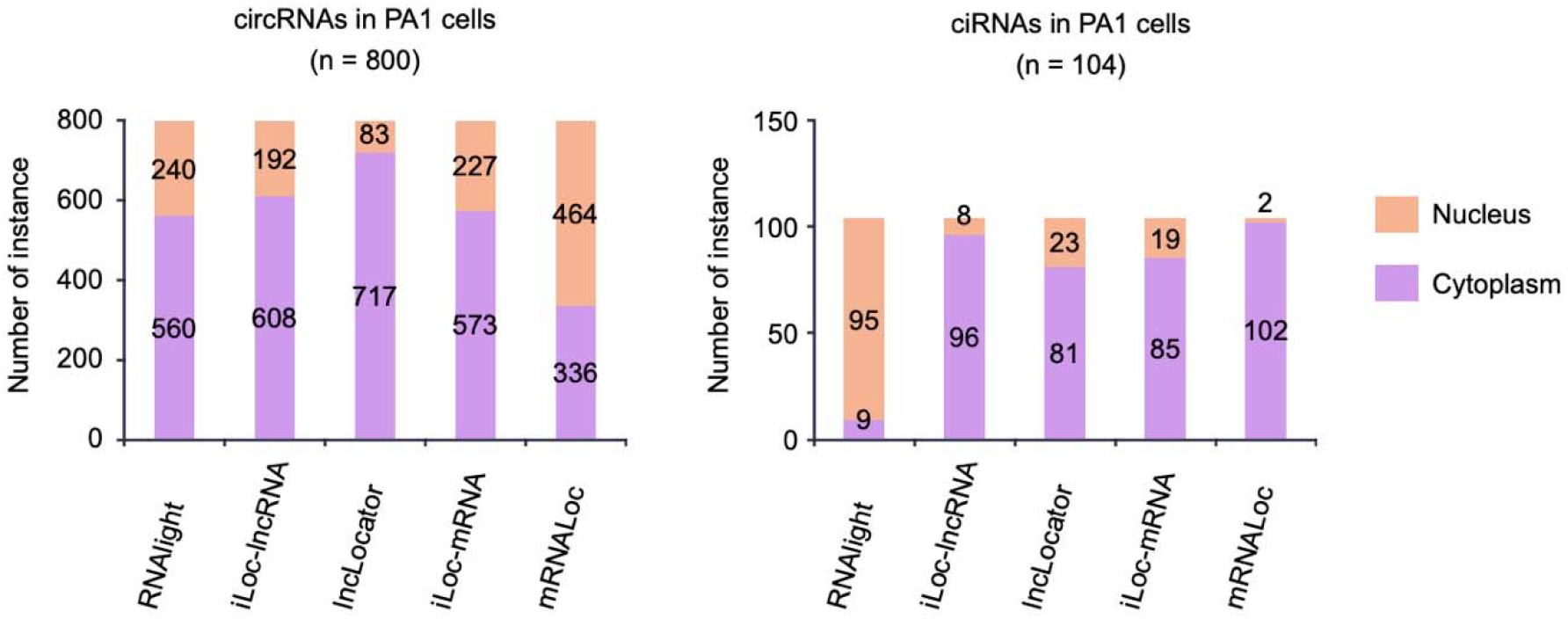
Benchmarking RNAlight and published models. RNAlight and other published models were compared by analyzing subcellular localizations of circular RNAs from back-spliced exons (circRNAs, left) or from spliced intron lariats (ciRNAs, right).

**Supplementary Figure S8.**
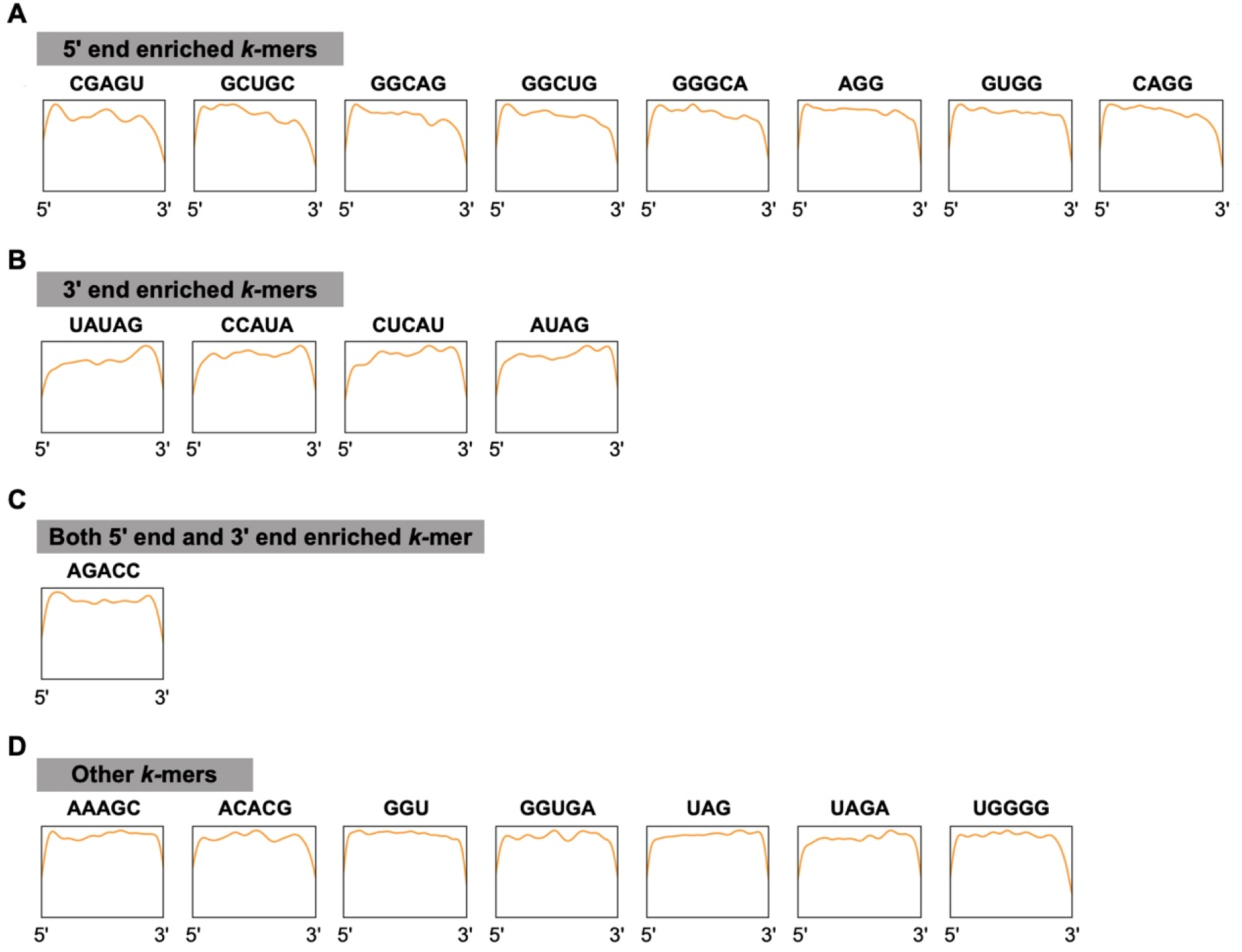
Analysis of lncRNA nucleus-related *k*-mer positional distribution. Positional distributions of 20 important lncRNA nucleus-related *k*-mers on nuclear lncRNAs (n = 1,986), which were classified to 5’ end enriched *k*-mers (**A**), 3’ end enriched *k*-mers (**B**), both 5’ end and 3’end enriched *k*-mer (**C**) and other *k*-mers (**D**).

**Supplementary Figure S9.**
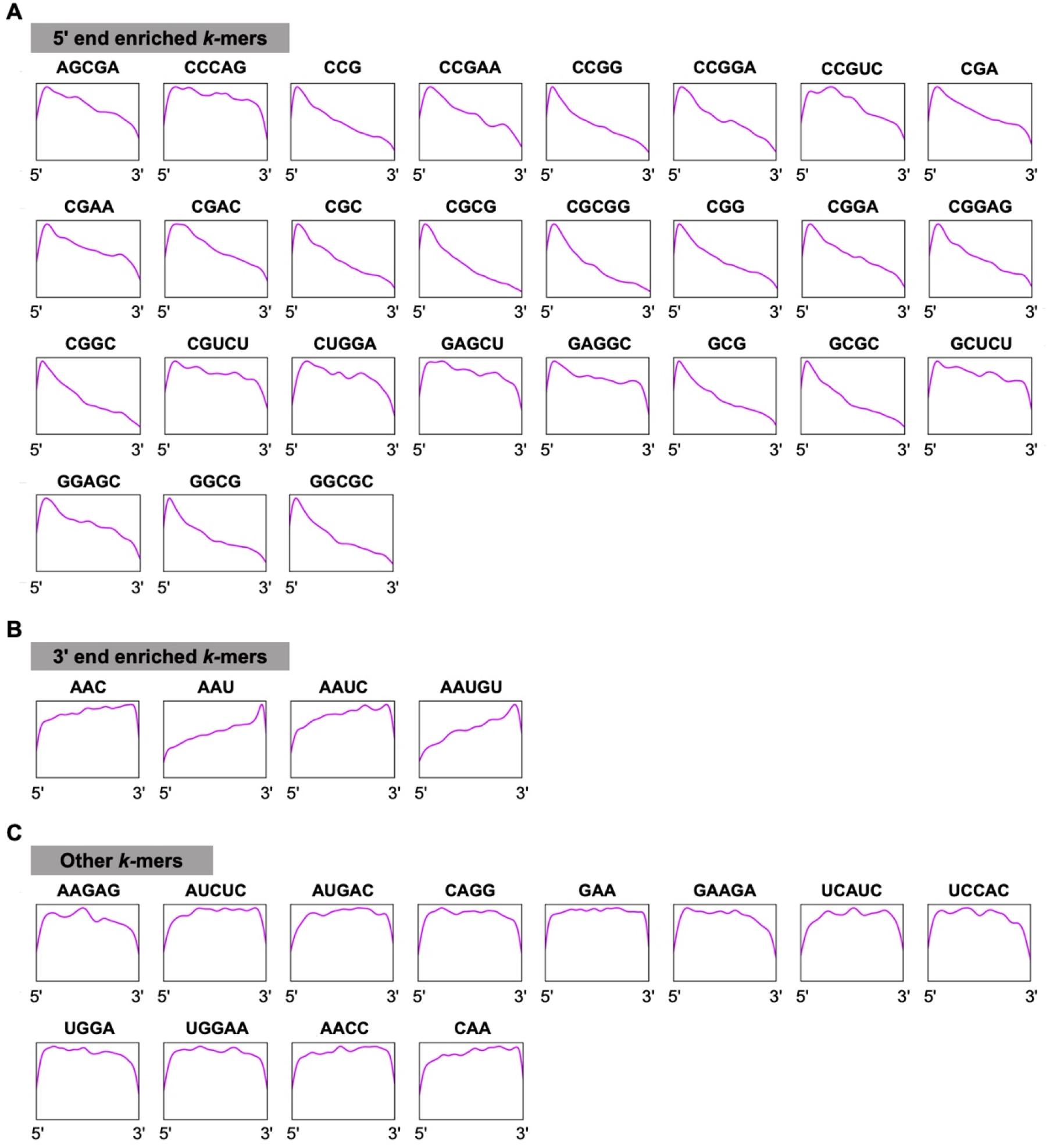
Analysis of lncRNA cytoplasm-related *k*-mer positional distribution. Positional distributions of 43 important lncRNA cytoplasm-related *k*-mers on cytoplasmic lncRNAs (n = 1,906), which were classified to 5’ end enriched *k*-mers (**A**), 3’ end enriched *k*-mers (**B**) and other *k*-mers (**C**).

**Supplementary Figure S10.**
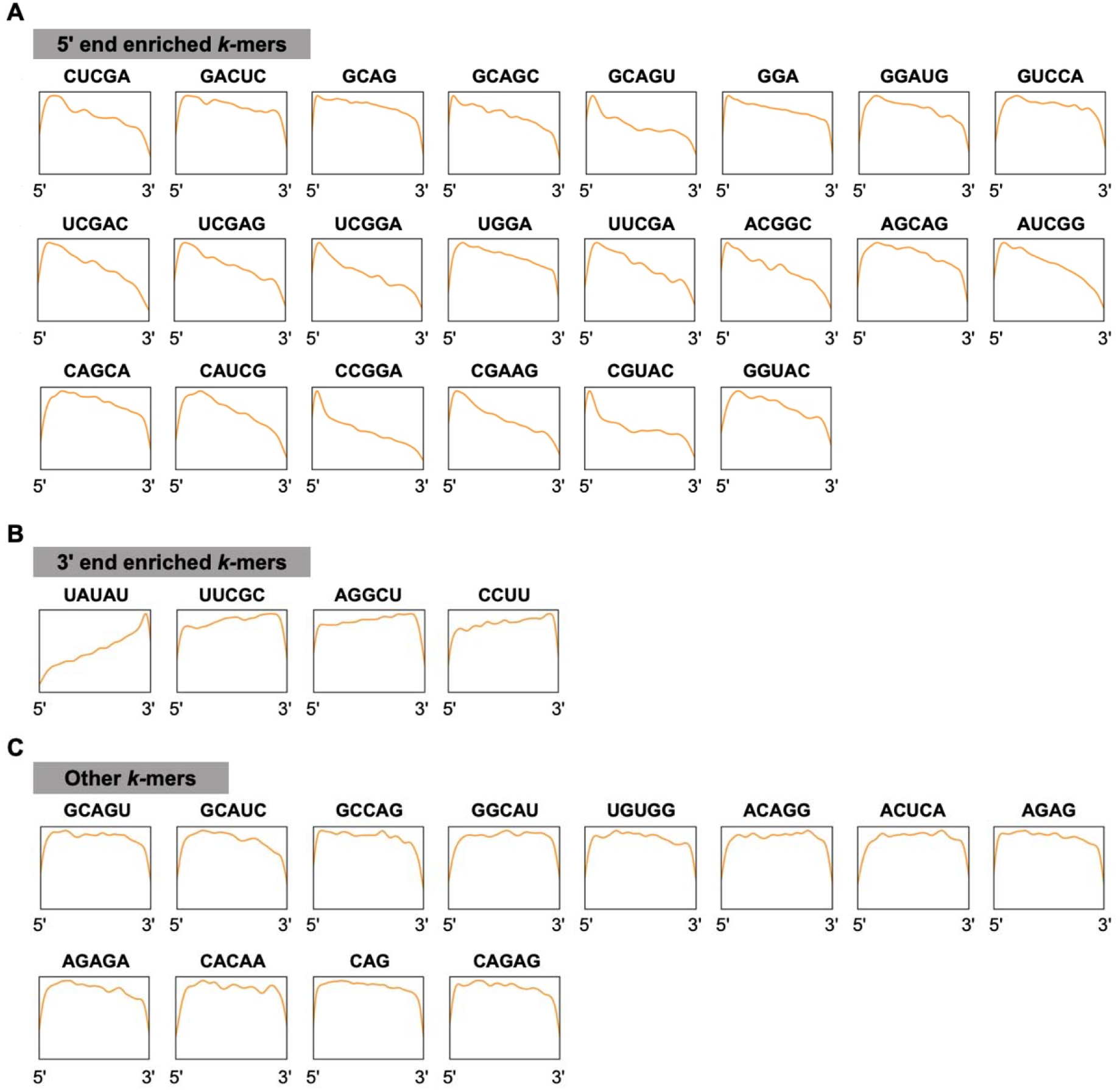
Analysis of mRNA nucleus-related *k*-mer positional distribution. Positional distributions of 38 important mRNA nucleus-related *k*-mers on nuclear mRNAs (n = 2,256), which were classified to 5’ end enriched *k*-mers (**A**), 3’ end enriched *k*-mers (**B**) and other *k*-mers (**C**).

**Supplementary Figure S11.**
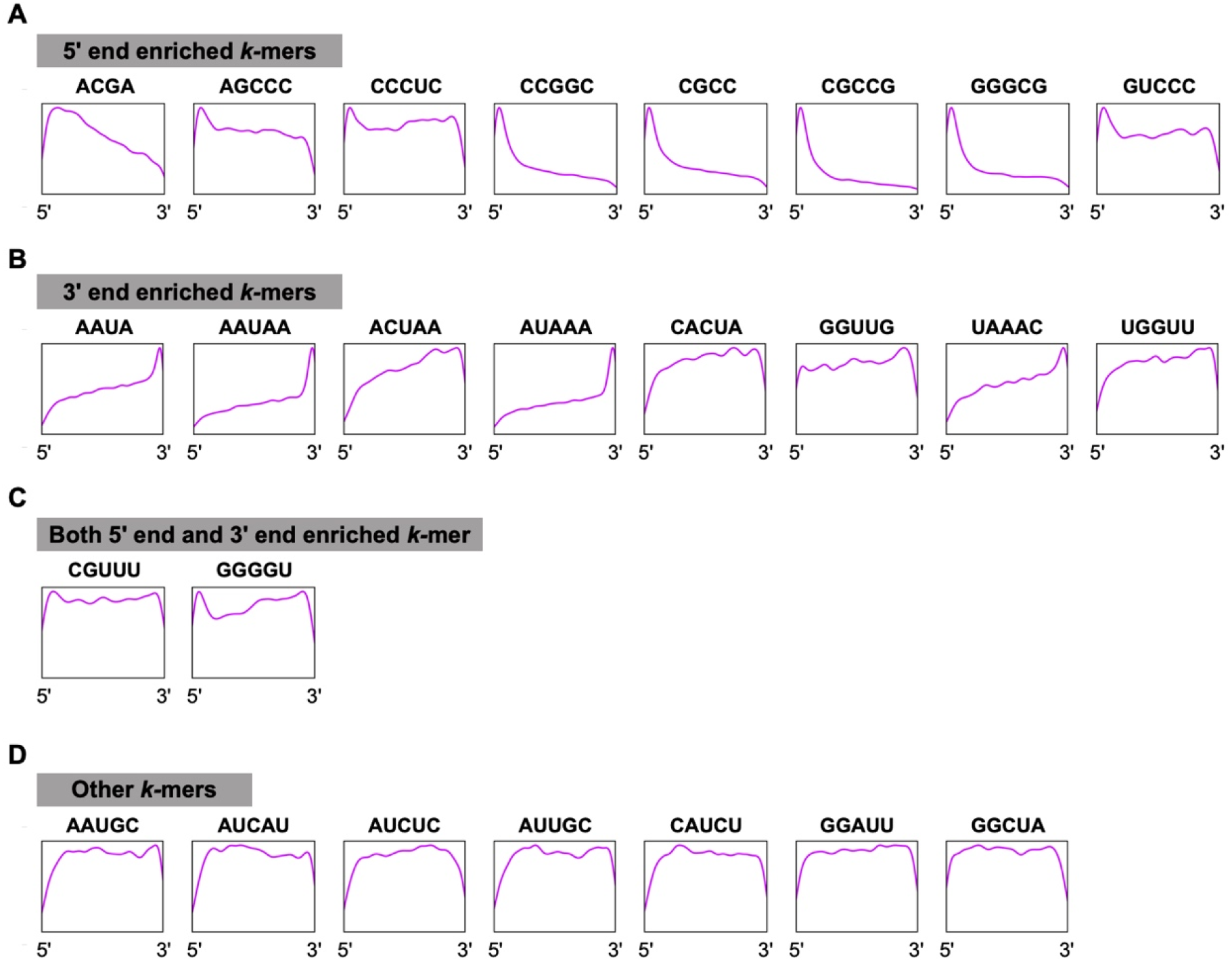
Analysis of mRNA cytoplasm-related *k*-mer positional distribution. Positional distributions of 25 important mRNA cytoplasm-related *k*-mers on cytoplasmic mRNAs (n = 2,924), which were classified to 5’ end enriched k-mers (**A**), 3’ end enriched *k*-mers (**B**), both 5’ end and 3’end enriched *k*-mers (**C**) and other *k*-mers (**D**).

**Supplementary Figure S12.**
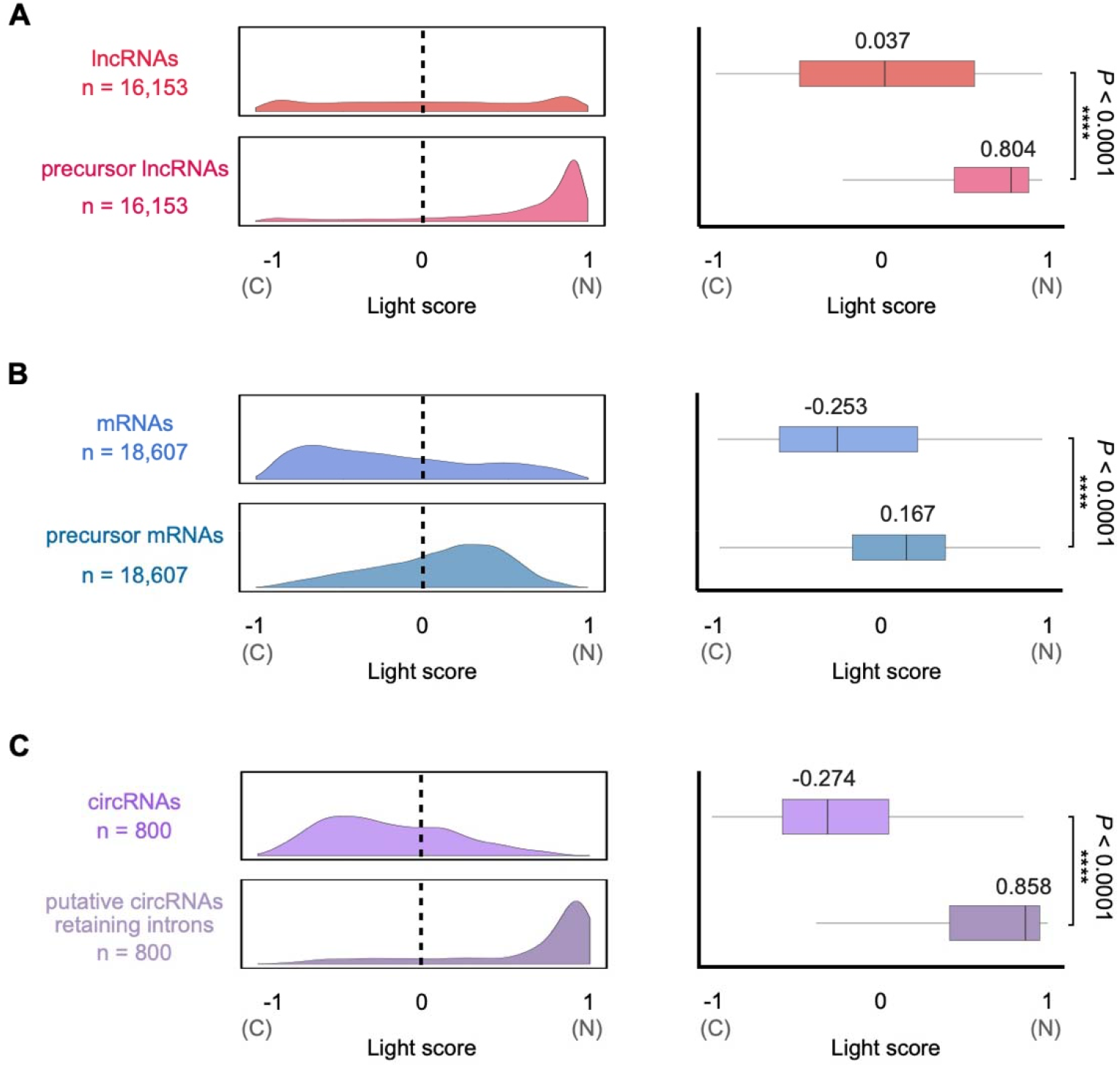
Application of RNAlight to predict subcellular localizations of various types of RNAs with/without intron sequence. Density (left) and box (right) charts show the distributions of Light scores reported by RNAlight across various types of RNAs with/without intron sequence, including lncRNAs (**A**) and mRNAs (**B**) from GENCODE v30 annotation, and circRNAs in PA1 cells from Zhang et al [61] (**C**). Statistical testing was performed with two-sided Welch’s *t*-test.

**Supplementary Figure S13.**
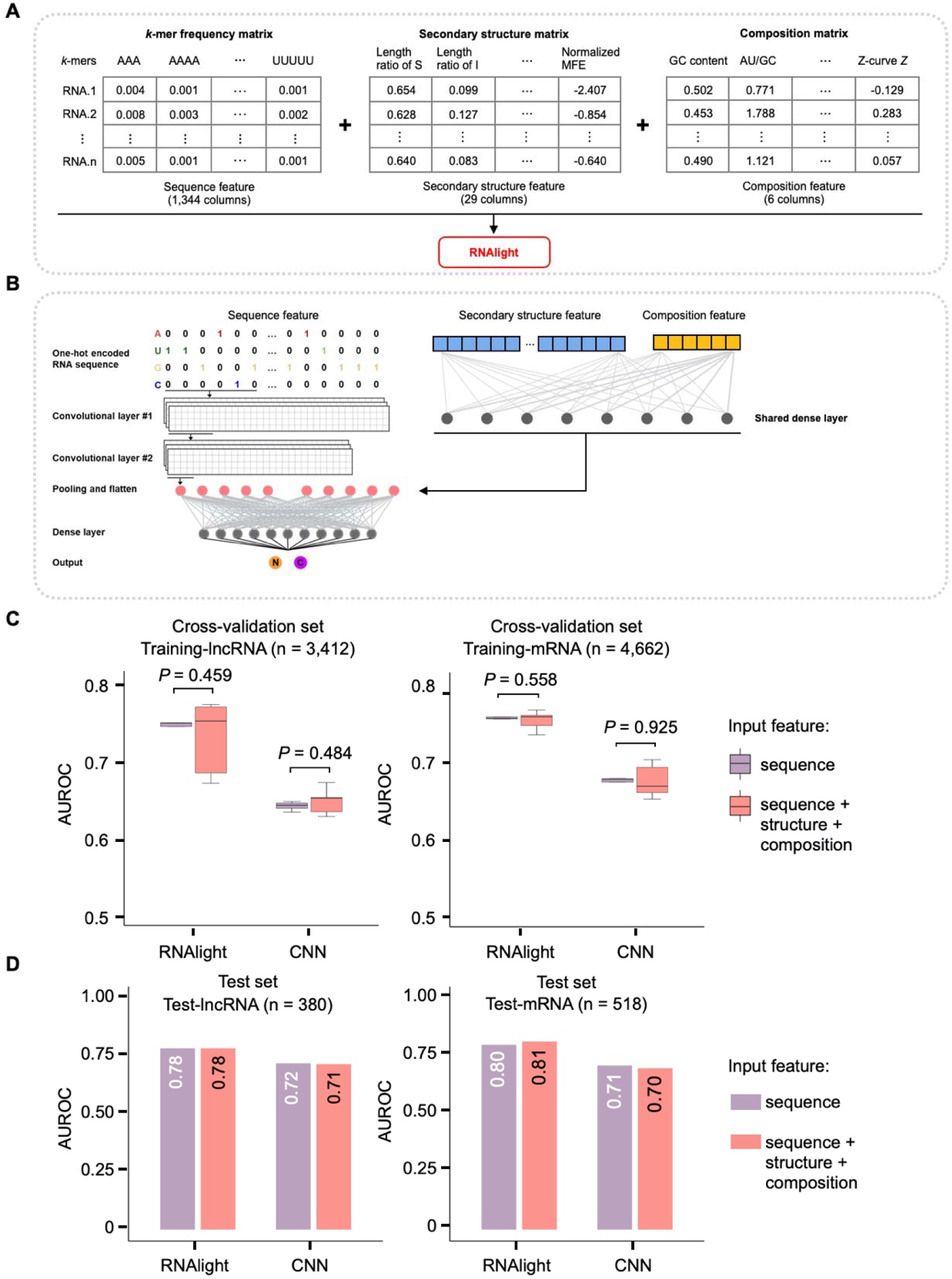
RNA subcellular localization prediction by *k*-mer frequency matrix with/without secondary structure and composition matrices. The *k*-mer frequency matrix, secondary structure matrix and composition matrix are combined to train machine learning (**A**) and deep learning (**B**) models (plum), which were shown with no significant improvement (**C, D**) compared to the training (salmon) when only using the *k*-mer frequency matrix.

## Supplementary table legends

**Supplementary Table S1. List of lncRNAs for model construction and evaluation in this study.** (A) List of lncRNAs for model training (Train-lncRNA). (B) List of lncRNAs for model testing (Test-lncRNA). Ensembl transcript ID, gene symbol, cDNA sequence and label are included.

**Supplementary Table S2. List of mRNAs for model construction and evaluation in this study.** (A) List of mRNAs for model training (Train-mRNA). (B) List of mRNAs for model testing (Test-mRNA). Ensembl transcript ID, gene symbol, cDNA sequence and label are included.

**Supplementary Table S3. An independent dataset from Halo-seq for model evaluation.** (A) LncRNAs from independent dataset of Halo-seq. (B) mRNAs from independent dataset of Halo-seq. Ensembl transcript ID, gene symbol, cDNA sequence and label are included.

**Supplementary Table S4. PCCs and *Z*-scores of 1,344 *k*-mers computed with the constructed lncRNA dataset from Supplementary Table S1.** *k*-mer, mean absolute SHAP value, PCC of *k*-mer frequency and SHAP value, *P*-value of PCC analysis, FDR of PCC analysis, *Z*-score of mean absolute SHAP value are listed.

**Supplementary Table S5. List of sequence features associated with lncRNA subcellular localization.** (A) List of lncRNA nucleus-related sequence features identified by *k*-mer assembling. (B) List of lncRNA cytoplasm-related sequence features identified by *k*-mer assembling. Sequence feature ID and sequence feature are included.

**Supplementary Table S6. List of sequence features matched to known RNA motifs and corresponding RBPs for lncRNA subcellular localization.** (A) List of lncRNA nucleus-related sequence features matched to known RNA motifs and corresponding RBPs. (B) List of lncRNA cytoplasm-related sequence features matched to known RNA motifs and corresponding RBPs. The list is shown by Query ID, Target ID, RBP, Optimal offset, *p*-value, *E*-value, *q*-value, Overlap, Query consensus, Target consensus and Orientation.

**Supplementary Table S7. PCCs and *Z*-scores of 1,344 *k*-mers computed with our mRNA dataset.** *k*-mer, mean absolute SHAP value, PCC of *k*-mer frequency and SHAP value, *P*-value of PCC analysis, FDR of PCC analysis, *Z*-score of mean absolute SHAP value are listed.

**Supplementary Table S8. List of sequence features associated with mRNA subcellular localization. (**A) List of mRNA nucleus-related sequence features identified by *k*-mer assembling. (B) List of mRNA cytoplasm-related sequence features identified by *k*-mer assembling. The list is shown by sequence feature ID and sequence feature.

**Supplementary Table S9. List of sequence features matched to known RNA motifs and corresponding RBPs for mRNA subcellular localization.** (A) List of mRNA nucleus-related sequence features matched to known RNA motifs and corresponding RBPs. (B) List of mRNA cytoplasm-related sequence features matched to known RNA motifs and corresponding RBPs. The list is shown by Query ID, Target ID, RBP, Optimal offset, *p*-value, *E*-value, *q*-value, Overlap, Query consensus, Target consensus and Orientation.

**Supplementary Table S10. Light scores of some specific RNAs with known subcellular localization.** RNA type, known localization, reference and Light score are listed.

